# Investigating Biases Associated with Dietary Starch Incorporation and Retention with an Oral Biofilm Model

**DOI:** 10.1101/2021.10.27.466104

**Authors:** Bjørn Peare Bartholdy, Amanda G. Henry

## Abstract

Dental calculus has proven to contain a wealth of information on the dietary habits of past populations. These insights have, to a large extent, been obtained by the extraction and identification of starch granules contained within the mineralised dental plaque from a wide range of regions and time periods. The scope of previous studies have been limited to microfossil extraction and identification to reconstruct dietary preferences from the archaeological record, and few studies have attempted to address the biases of starch retention in dental calculus. Those that have considered this problem have been limited to *in vivo* studies on modern humans and non-human primates. Here, we present a multispecies oral biofilm model, which allows experimental research on starch incorporation and retention to be conducted on *in vitro* dental calculus in a controlled laboratory setting. The biofilms were exposed to treatment solutions with known quantities of dietary starches (wheat and potato) during the 25-day growth period. After this, the starch granules were extracted from the mature biofilm (by dissolution in EDTA), and counted. We show that the granule counts extracted from the model dental calculus represented a low proportion (ranging from 0.06% to 0.16%) of the total number of granules exposed to the biofilms throughout the experiment. Additionally, we found that the ratios of granule sizes from the extracted starch granules differed from the original treatment solutions, with large granules (>20 *μ*m) consistently being under-represented. We also found a correlation between the absolute granule counts and dry-weight of the biofilm (*r* = 0.66, 90%CI[0.46,0.79]), as well as between the concentration (count per mg) of granules and dry-weight (*r* = 0.30, 90%CI[0.06,0.51]).

Our results reinforce previous *in vivo* studies suggesting that dental calculus presents a very small, and partly biased picture of the original dietary intake of starches, with an over-representation of plants producing granules smaller than 20 *μ*m in size. The experimental model presented here is well-suited to address the need for further validation of methods and biases associated with dietary research on dental calculus.

## 1 Introduction

Dental calculus has proven to contain a wealth of dietary information in the form of plant microfossils (Hardy et al., 2009; Henry & Piperno, 2008), proteins (Hendy et al., 2018; Warinner, Hendy, et al., 2014), and other organic residues (Buckley et al., 2014). This dietary information can be preserved within the mineralised dental plaque over many millennia, providing a unique window into the food-related behaviours of past populations (Henry & Piperno, 2008; Jovanović et al., 2021; Tao et al., 2020) and extinct species (Hardy et al., 2012; Henry et al., 2014).

Until recently, only a few studies directly investigated the presence of plant microremains in the dental calculus of archaeological remains. The ability to extract phytoliths from the dental calculus of archaeological fauna to investigate diet was first noted by Armitage (1975), and later by Middleton and Rovner (1994), and Fox and colleagues (1996). Starches and phytoliths were extracted from human dental calculus by Cummings and Magennis (1997).

In more recent years, the study of dental calculus has increased exponentially, and the wealth of information contained within the mineralised matrix has largely been acknowledged. The use of dental calculus spans a wide variety of archaeological research areas, such as oral microbiome characterisation (including pathogens) through the analysis of DNA and proteins (Adler et al., 2013; Warinner, Rodrigues, et al., 2014), microbotanical remains (Hardy et al., 2009; Henry & Piperno, 2008; Mickleburgh & Pagán-Jiménez, 2012), other organic residues and proteins from dietary compounds (Buckley et al., 2014; Hendy et al., 2018), and nicotine use (Eerkens et al., 2018). Especially the extraction of starch granules has become a rich source of dietary information, as starch granules have proven to preserve well within dental calculus over a variety of geographical and temporal ranges (Henry et al., 2014; Jovanović et al., 2021; Piperno & Dillehay, 2008; Tao et al., 2020).

Despite this, our knowledge of dental calculus and the incorporation pathways of the various markers is limited (Radini et al., 2017), as is our knowledge of information-loss caused by these pathways. Additionally, the methods we use to extract and analyse dental calculus, and make inferences on past diets represent another potential source of bias. Studies on both archaeological and modern individuals have explored these biases in more detail. Extraction methods were tested by Tromp and colleagues (2017), specifically regarding decalcification using HCl or EDTA. The authors found significantly more starches with the EDTA extraction method than the HCl extraction method; however, as noted by the authors, comparisons involving archaeological calculus are problematic due to variability between and within individuals. Studies conducted on modern humans (Leonard et al., 2015) and non-human primates (Power et al., 2015), have explored how well microremains (phytoliths and starches) extracted from dental calculus represent the actual dietary intake. These studies are justifiably limited, despite meticulous documentation and observation, due to unknown variables and uncertainty involved when studying living organisms. Dental calculus is a complex oral biofilm with a multifactorial aetiology and variable formation rates both within and between individuals (Haffajee et al., 2009; Jepsen et al., 2011), contributing to the stochasticity of starch representation being observed in numerous studies. Additionally, the concentration of oral *α*-amylase differs both between and within individuals (Froehlich et al., 1987; Nater et al., 2005), causing different rates of hydrolysis of the starch granules present in the oral cavity. Add to this the effects of the many different methods of starch processing, as well as post-depositional processes that are still being explored (García-Granero, 2020), and it becomes clear that using dental calculus to reconstruct diet is a highly unpredictable process.

In this exploratory study, we use an oral biofilm model to investigate the retention of starch granules within dental calculus in a controlled laboratory setting, allowing us full control over dietary input. Our main questions concern the representation of granules extracted from the calculus compared to the actual intake. How much of the original diet is incorporated into the calculus, and how much is recovered? Is there differential loss of information from specific dietary markers that affects the obtained dietary information, and how does this affect the representation of diet from extracted microremains?

We find that, despite the absence of *α*-amylase in the model, a limited proportion of the starch input is actually retained in the calculus. We also observed a shift in the size ratios of individual starch granules that are incorporated into the calculus, and that the number of incorporated starch granules increases as the size of the calculus deposit increases.

## 2 Materials and Methods

### 2.1 Biofilm formation

In this study we employ a multispecies oral biofilm model following a modified protocol from Sissons and colleagues (1991) and Shellis (1978). In brief, a biofilm inoculated with whole saliva was grown on a substrate suspended in artificial saliva, and fed with sugar (sucrose). After several days of growth, the biofilm was exposed to starch solutions. Mineralisation of the biofilm was aided by exposure to a calcium phosphate solution. After 25 days of growth, the mineralised biofilm was collected for further analysis. The setup comprises a polypropylene 24 deepwell PCR plate (KingFisher 97003510) with a lid containing 24 pegs, which is autoclaved at 120°C, 1 bar overpressure, for 20 mins. The individual pegs were the substrata on which the calculus grew. Using this system allowed for easy transfer of the growing biofilm between saliva, feeding solutions, and mineral solutions.

The artificial saliva (AS) is a modified version of the basal medium mucin (BMM) described by Sissons and colleagues (1991). It contains 2.5 g/l partially purified mucin from porcine stomach (Type III, Sigma M1778), 5 g/l trypticase peptone (Roth 2363.1), 10 g/l proteose peptone (Oxoid LP0085), 5 g/l yeast extract (BD 211921), 2.5 g/l KCl, 0.35 g/l NaCl, 1.8 mmol/l CaCl_2_, 5.2 mmol/l Na_2_HPO_4_ (Sissons et al., 1991), 6.4 mmol/l NaHCO_3_ (Shellis, 1978), 2.5 mg/l haemin. This is subsequently adjusted to pH 7 with NaOH pellets and stirring, autoclaved (15 min, 120°C, 1 bar overpressure), and supplemented with 5.8 *μ*mol/l menadione, 5 mmol/l urea, and 1 mmol/l arginine (Sissons et al., 1991).

Fresh whole saliva (WS) for inoculation was provided by a 31-year-old male donor with no history of caries, who abstained from oral hygiene for 24 hours. No food was consumed two hours prior to donation and no antibiotics were taken up to six months prior to donation. The saliva was filtered through a sterilised (with sodium hypochlorite, 10–15% active chlorine) nylon cloth to remove particulates. Substrata were inoculated with 1 ml/well of a two-fold dilution of WS in sterilised 20% (v/v) glycerine for four hours at 36°C, to allow attachment of the salivary pellicle and plaque-forming bacteria. After initial inoculation, the substrata were transferred to a new plate containing 1 ml/well AS and incubated in a shaking incubator (Infors HT Ecotron) at 36°C, 30 rpm. The inoculation process was repeated on days 3 and 5. AS was partially refreshed once per day and fully refreshed every three days, throughout the experiment, by transferring the substrata to a new plate containing stock AS. To feed the bacteria, the substrata were transferred to a new plate, containing 5% (w/v) sucrose, for six minutes twice daily, except on inoculation days (days 0, 3, and 5), where the samples only received one sucrose treatment after inoculation.

Starch treatments were initiated on day 9 to avoid starch granule counts being affected by *α*-amylase hydrolysis from saliva inoculation days. An *α*-amylase (EC 3.2.1.1) activity assay was conducted to confirm that no amylase was present in the system before starch treatments started. Starch treatments replaced sucrose treatments, occurring twice per day for six minutes. The starch treatments involved transferring the substrata to a new plate containing a 0.25% (w/v) starch from potato (Roth 9441.1) solution, a 0.25% (w/v) starch from wheat (Sigma S5127) solution, and a 0.5% (w/v) mixture of equal concentrations (w/v) wheat and potato. All starch treatments were created in dH_2_O with 5% (w/v) sucrose. Before transferring biofilm samples to the starch treatment plate, the plates were agitated to keep the starches in suspension in the solutions. During treatments, the rpm was increased to 60 to facilitate contact between starch granules and biofilms.

After 15 days, mineralisation was encouraged with a calcium phosphate monofluorophosphate urea (CPMU) solution containing 20 mmol/l CaCl_2_, 12 mmol/l NaH_2_PO_4_, 5 mmol/l Na_2_PO_3_F, 500 mmol/l urea, and (0.04 g/l MgCl) (Pearce & Sissons, 1987; Sissons et al., 1991). The substrata were submerged in 1 ml/well CPMU for six minutes, five times daily, in a two-hour cycle. During the mineralisation period, starch treatments were reduced to once per day after the five CPMU treatments. This process was repeated for 10 days until the end of the experiment on day 24 (see Figure 1 for an overview of the protocol).

**Figure 1:**
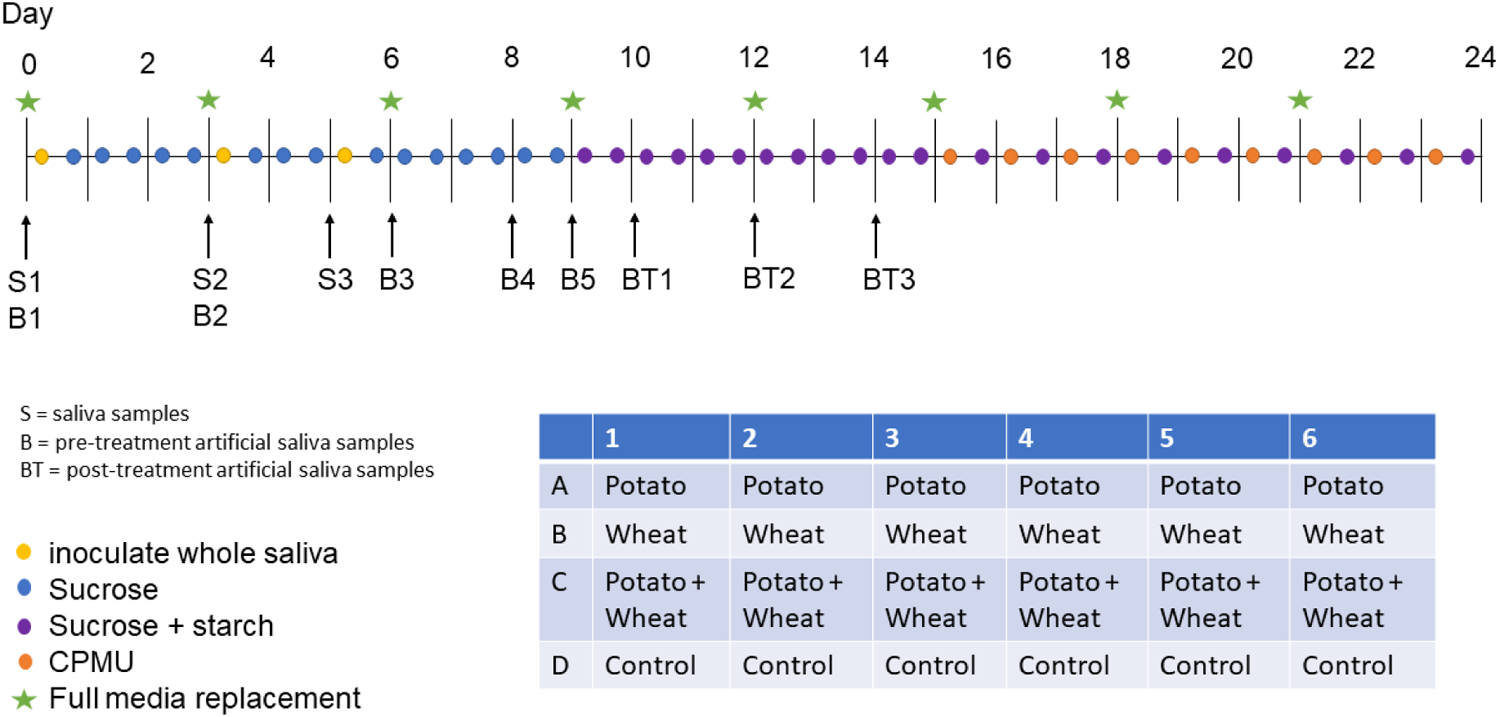
Overview of experiment protocol including the plate setup.

All laboratory work was conducted in sterile conditions under a laminar flow hood to prevent starch and bacterial contamination. Control samples that only received sucrose as a treatment were included to detect starch contamination from the environment or cross-contamination from other wells in the same plate.

### 2.2 Amylase activity detection

An *α*-amylase (EC 3.2.1.1) activity assay was conducted on artificial saliva samples collected from the plate wells on days 3, 6, 8, 9, 10, 12, and 14. Whole saliva samples were collected on days 0, 3, and 5 as positive controls. Collected samples were stored at 4°C until the assay was conducted on day 18. All samples and standard curves were run in triplicates on two separate plates. Positive control saliva samples were compared against a standard curve containing H_2_O, while artificial saliva samples were compared against a standard curve containing sterile artificial saliva (due to the colour of artificial saliva). Two photometric readings were conducted for each plate with a 540 nm filter on a Multiskan FC Microplate Photometer (Thermo Scientific 51119000). The protocol is a slightly modified version of an Enzymatic Assay of *α*-Amylase (https://www.sigmaaldrich.com/NL/en/technical-documents/protocol/protein-biology/enzyme-activity-assays/enzymatic-assay-of-a-amylase) (Bernfeld, 1955), which measures the amount of maltose released from starch by *α*-amylase activity. Results are reported in units (U) per mL enzyme, where 1 U releases 1 mg of maltose in 6 minutes.

### 2.3 Treatment solutions

A 1 ml aliquot of each starch solution was taken, from which 10 *μ*l was mounted on a microscope slide with an 18 × 18 mm coverslip, and counted under a light microscope (Zeiss Axioscope A1). For wheat and mixed treatment samples, we counted three slide transects (at ca. 1/4, 1/2, and 3/4 of the slide), and the sample counts were extrapolated to the total number of granules exposed to the samples over 16 days of treatments (see Supplementary Material for more details). For potato treatment samples, the whole slide was counted.

### 2.4 Extraction method

Extraction of starches from the calculus samples was performed by dissolving the calculus in 0.5 *M* ethylenediaminetetraacetic acid (EDTA) (Le Moyne & Crowther, 2021; Modi et al., 2020; Tromp et al., 2017), and vortexing for 3 days until the sample was completely dissolved. Twenty *μ*l of sample was mounted onto a slide with an 18×18 mm coverslip. When transferring the sample to the slide, the sample was homogenised using the pipette to ensure that the counted transects were representative of the whole slide. The count from the slide was extrapolated to the whole sample (see Supplementary Material for more detail).

Both wheat and potato granules were divided into three size categories: small (<10 *μ*m), medium (10 – 20 *μ*m), and large (>20 *μ*m).

### 2.5 Statistical analysis

Statistical analysis was conducted in R version 4.1.1 (2021-08-10) (R Core Team, 2020) and the following packages: tidyverse (Wickham et al., 2019), broom (Robinson et al., 2021), here (Müller, 2020), and patchwork (Pedersen, 2020).

To see if biofilm growth was differently affected by starch treatments, a one-way ANOVA with sample weight as the dependent variable (DV) and treatment as the grouping variable (GV) was conducted. To analyse granule counts and calculate size proportions, mean counts for each treatment were taken across both experimental plates, resulting in a mean count for each granule size category within each treatment.

Pearson’s *r* was conducted on sample weight and total starch count, as well as sample weight and starch count per mg calculus. The total count for each sample within a treatment was standardised by z-score to account for the differences in magnitude between the potato and wheat counts. This was applied to total biofilm weight and starch count per mg calculus (also z-score standardised) to account for differences in starch concentration in the calculus (as per Wesolowski et al., 2010).

## 3 Results

All samples yielded su?icient biofilm growth and starch incorporation to be included in the analysis (Figure 2), resulting in a total of 48 biofilm samples (two plates of 24), 45 of which were used for analysis (three samples were set aside for later analysis). Most control samples contained no starch granules, while some contained negligible quantities (see Supplementary Material).

**Figure 2:**
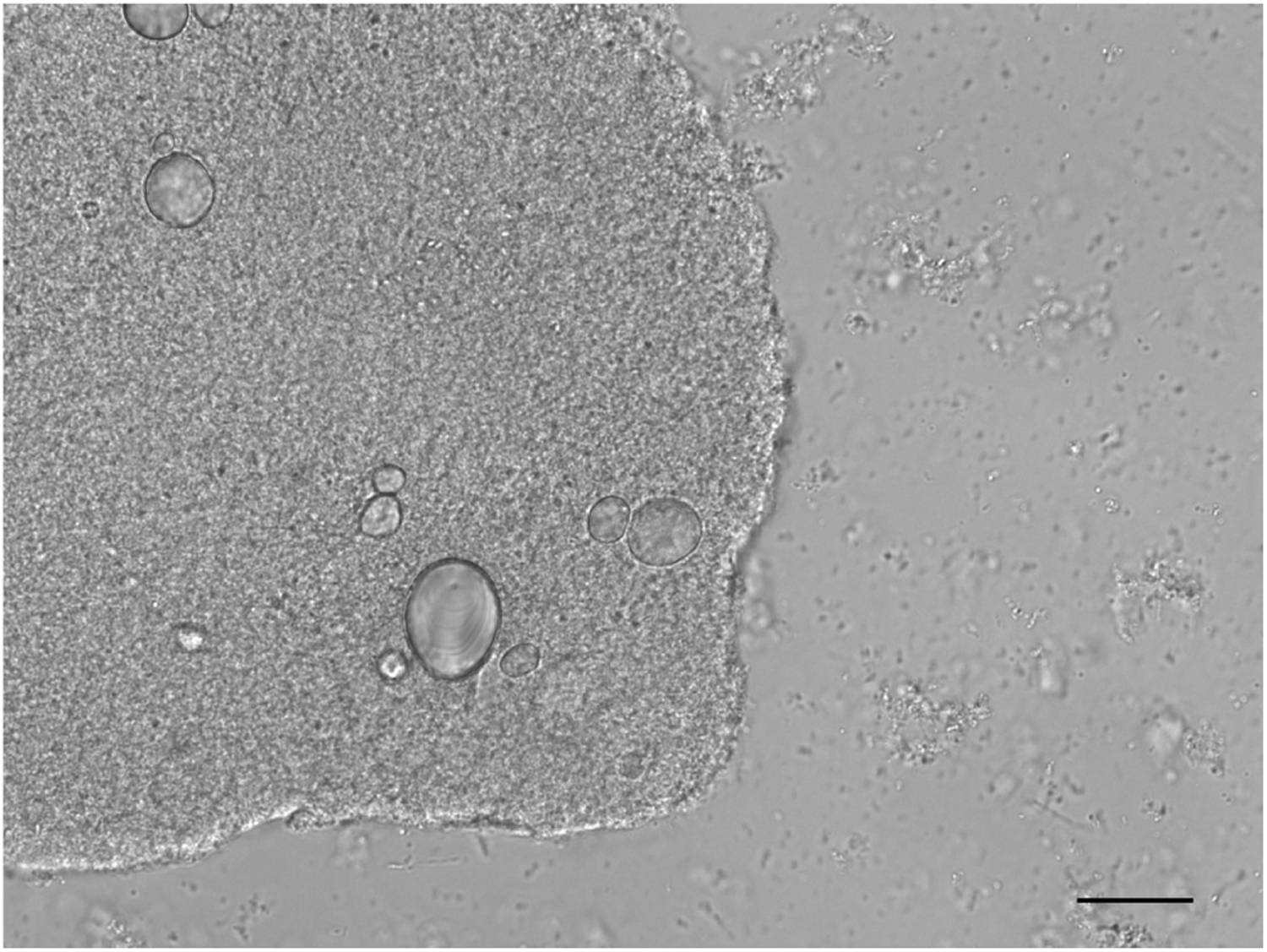
Microscope image of a biofilm sample exposed to the potato starch solution. Potato granules can be seen within a bacterial community. Scale bar = 20 *μ*m.

### 3.1 No amylase activity detected in the model

No *α*-amylase activity was detected in any of the artificial saliva samples from any of the days that were sampled. Only positive controls contained amylase activity that could be detected in the assay, ranging from 3.4 to 10.3 U/mL enzyme (full results can be found in the Supplementary Material). The results are not comparable to other studies presenting *α*-amylase activity levels in humans, as the unit definition may differ; however, they are su?icient to show that there is no activity in the system.

### 3.2 Treatment type had minimal effect on biofilm growth

A one-way ANOVA suggests that the type of starch used during the biofilm growth period had a minimal effect on the growth of the biofilm (expressed as total dry weight of the sample), F(3, 43) = 1.16, p = 0.335. A summary of sample weights is available in Table 1.

**Table 1:** Summary statistics for biofilm dry-weights (in mg) by treatment.

| Treatment | Mean | SD | Min | Max |
| --- | --- | --- | --- | --- |
| control | 5.44 | 2.45 | 1.67 | 11.20 |
| mix | 4.28 | 1.95 | 1.50 | 8.44 |
| potato | 6.25 | 2.07 | 2.54 | 8.92 |
| wheat | 5.53 | 3.45 | 0.56 | 9.80 |

### 3.3 Starch counts

It was not possible to differentiate between potato and wheat starches smaller than ca. 10 *μ*m. These were counted as wheat, as we assumed that the majority of the small granules were wheat. We make this assumption based on the counts of small starches in the wheat-only and potato-only solutions. Of the total amount of small starches in these two solutions, 99.2% are from wheat.

The separate wheat and potato solutions were made with a 0.25% (w/v) starch concentration, while the mixed-starch solution was made with 0.25% (w/v) of each starch, with a total concentration of 0.50% (w/v). The mixed treatment had the highest absolute count of starch granules in solution (mean = 2.9 × 10^7^), while the biofilms exposed to the wheat solution preserved the greatest number of granules (mean = 2.77 × 10^4^). The potato treatment had the lowest absolute counts in both the solution (3.02 × 10^6^) and in the biofilm samples (4850) (Tables 2 and 3).

**Table 2:** Mean starch counts from solutions, including the proportional makeup of the different sizes of granules.

| Solution | Starch | Small (%) | Medium (%) | Large (%) | Total (%) |
| --- | --- | --- | --- | --- | --- |
| mix | potato |  | 1051733 (53.1%) | 928000 (46.9%) | 1979733 (100.0%) |
| mix | wheat | 18838400 (69.7%) | 6403200 (23.7%) | 1794133 (6.6%) | 27035733 (100.0%) |
| mix | both | 18838400 (64.9%) | 7454933 (25.7%) | 2722133 (9.4%) | 29015467 (100.0%) |
| potato | potato | 123733 (4.1%) | 1337867 (44.4%) | 1554400 (51.5%) | 3016000 (100.0%) |
| wheat | wheat | 16139467 (63.5%) | 6434133 (25.3%) | 2830400 (11.1%) | 25404000 (100.0%) |

**Table 3:** Mean starch counts extracted from samples with standard deviation (SD), including the proportion of granule sizes of the total count.

| Treatment | Starch | Small (%) | SD | Medium (%) | SD | Large (%) | SD | Total (%) | SD |
| --- | --- | --- | --- | --- | --- | --- | --- | --- | --- |
| mix | potato |  |  | 1959 (79.6%) | 1801 | 501 (20.40%) | 446 | 2460 (100%) | 2189 |
| mix | wheat | 9515 (54.60%) | 8860 | 6522 (37.4%) | 6026 | 1381 (7.93%) | 1196 | 17417 (100%) | 15878 |
| mix | both | 9515 (47.90%) | 8860 | 8480 (42.7%) | 7653 | 1882 (9.47%) | 1596 | 19877 (100%) | 17768 |
| potato | potato | 351 (7.24%) | 297 | 3565 (73.6%) | 2402 | 930 (19.20%) | 929 | 4846 (100%) | 3316 |
| wheat | wheat | 15235 (55.00%) | 11944 | 12148 (43.9%) | 11052 | 1953 (7.06%) | 2016 | 27680 (100%) | 23554 |

#### 3.3.1 Proportion of available starches incorporated in samples

The proportion of total starches from the solutions that were incorporated into the samples ranged from 0.06% to 0.16%, with potato granules being more readily incorporated than wheat in both the separated- and mixed-treatment samples (Table 4). There is an inverse relationship between the absolute starch count in the solutions and the proportional incorporation of starches in the biofilm samples, i.e., potato had the lowest absolute count in solutions, but the highest proportional incorporation, and vice versa for the mixed treatment.

**Table 4:** The mean percentage of starches from the solutions that were incorporated into the samples.

| Treatment | Starch | Small | Medium | Large | Total |
| --- | --- | --- | --- | --- | --- |
| mix | potato |  | 0.19% | 0.05% | 0.12% |
| mix | wheat | 0.05% | 0.10% | 0.08% | 0.06% |
| mix | both | 0.05% | 0.11% | 0.07% | 0.07% |
| potato | potato | 0.28% | 0.27% | 0.06% | 0.16% |
| wheat | wheat | 0.09% | 0.19% | 0.07% | 0.12% |

Wheat incorporation was most affected in the mixed-treatment samples, with only 0.06% of the total available starches being incorporated into the sample, compared to 0.16% in the separated wheat treatment.

#### 3.3.2 Size ratios differ between solutions and samples

Overall, medium starch granules had a higher mean rate of incorporation (0.171%) than small (0.120%) and large (0.066%) starch granules across all treatments, while large potato starches had the lowest rate of incorporation across all treatments.

The difference in incorporation between the size categories resulted in a change in size ratios between the original starch solutions and the extracted samples. Large potato granules (> 20 *μ*m) were most affected, with a 32.3% decrease in relative abundance in the potato-only treatment, and a 26.5% decrease in mixed treatments. Medium granules increased in relative abundance across all samples, while small granules decreased in wheat treatments and increased in potato treatments (Figure 3).

**Figure 3:**
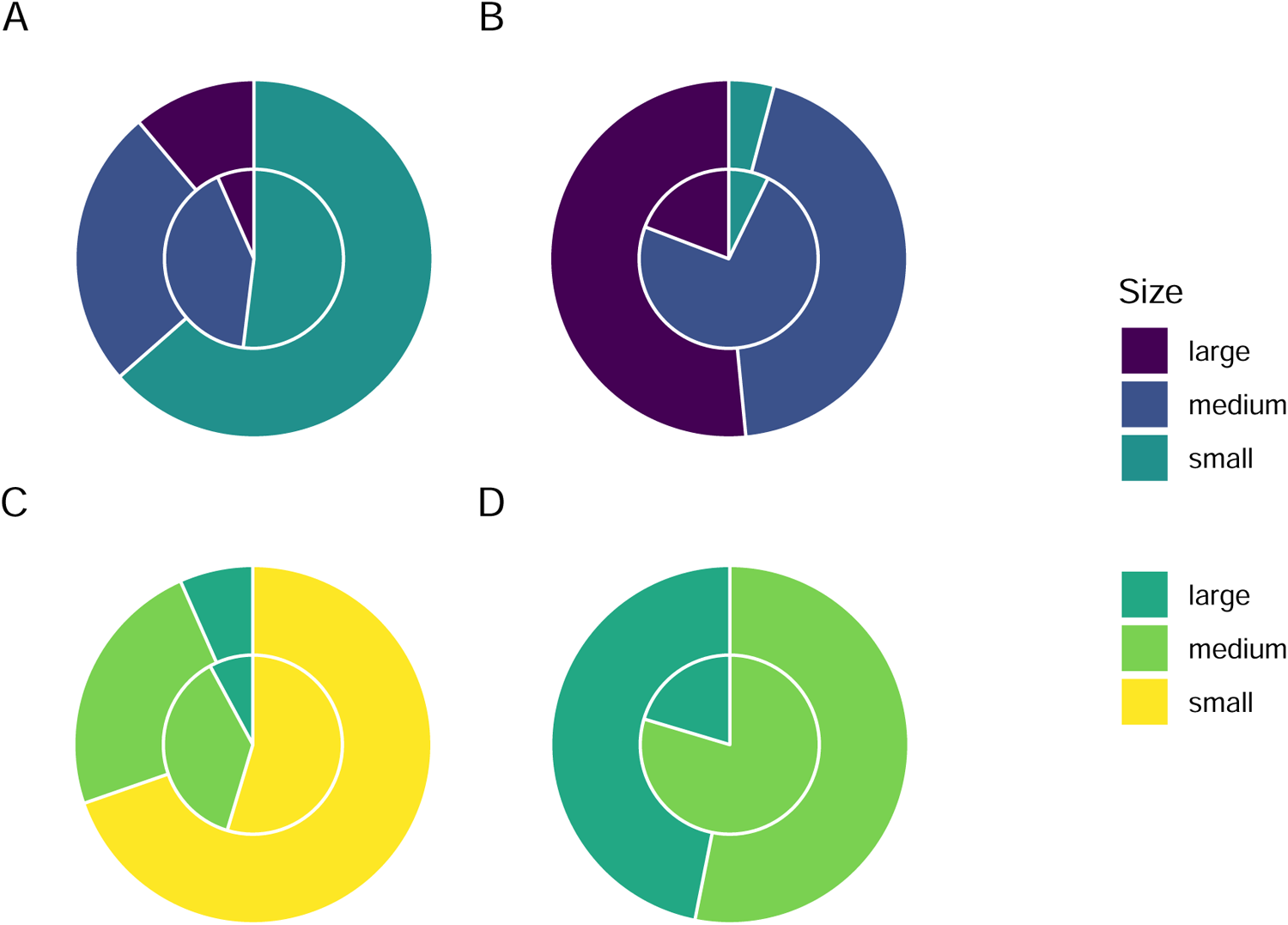
Proportion of sizes of starch granules from solutions (outer ring) and treatment samples (inner ring) in separated wheat (A) and potato (B) treatments, and mixed wheat (C) and potato (D) treatments.

#### 3.3.3 Biofilm weight correlated positively with extracted starch counts

Pearson’s *r* suggests a strong positive correlation between the total weight of the biofilms and the total starch count (standardised by z-score) extracted from the samples across treatments, *r* = 0.659, 90%CI[0.463, 0.794], p < 0.001 (Figure 4A).

**Figure 4:**
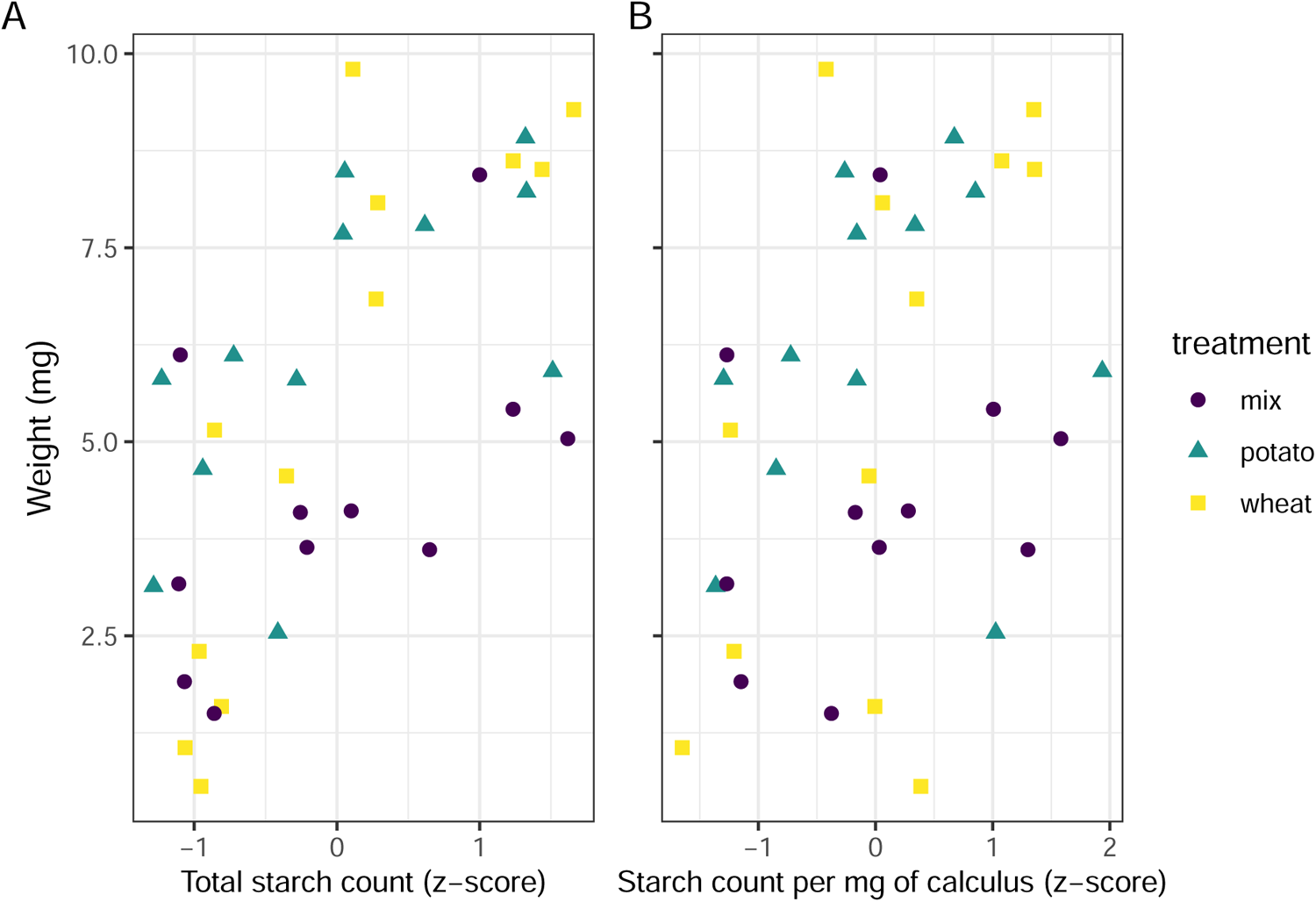
Scatter plots of (A) sample weight in mg and standardised starch count by z-score for separated treatments, and (B) sample weight in mg and standardised count of starch grains per mg calculus.

The same test was applied to total biofilm weight and starch count per mg calculus (also standardised by z-score), resulting in a weak positive correlation, *r* = 0.3, 90%CI[0.0618, 0.506], p = 0.0403 (Figure 4B).

## 4 Discussion

Here, we have provided a method for exploring the incorporation of dietary starches into the mineral matrix of a dental calculus biofilm model. Our results show that a very low proportion of the starches exposed to the biofilm during growth are retained in the mineral matrix, and that the size of the starch granules may affect the likelihood of incorporation. The proportions of starch granules (of all sizes) present in the extracted samples were similar across all treatments (0.06% to 0.16%), despite large differences in absolute granule counts between wheat (mean = 25,404,000) and potato (mean = 3,016,000) solutions. The absolute counts, however, differed more visibly between treatments and was proportional with the total count of granules in the treatment solutions. Wheat and mixed solutions had the highest absolute mean count of starch granules, and also had the highest absolute mean count of starch granules extracted from the dental calculus (Tables 2 and 3). This suggests that the starches that are more frequently consumed will be present in higher quantities in the dental calculus, at least prior to inhumation and degradation in the burial environment. Despite the low proportion of granules recovered from the model calculus (0.06% to 0.16%), the absolute counts were still substantially greater than counts recovered from archaeological remains (Tromp et al., 2017; Tromp & Dudgeon, 2015; Wesolowski et al., 2010), which could in part be due to the lack of oral amylase activity in our model.

We have also shown that the size of the starch granules influences the likelihood of incorporation into the calculus. Starch granules larger than 20 *μ*m in maximum length were underrepresented in the calculus samples compared to the original starch solutions, an effect that was consistent across all three treatments, while medium granules (10–20 *μ*m) were often over-represented (Table 4, and Figure 3). Large potato granules were most affected, potentially because of the greater size-range. They can reach up to 100 *μ*m in maximum length, whereas wheat granules generally only reach up to 35 *μ*m (Gismondi et al., 2019; Haslam, 2004; Seidemann, 1966, pp. 174–176). Granule morphology may also play a role. Large wheat granules are lenticular and have a larger surface area compared to volume, whereas large potato granules are ovoid and have a larger volume compared to surface area (Jane et al., 1994; Reichert, 1913, pp. 364–365; Seidemann, 1966, pp. 174–176; van de Velde et al., 2002). Another potentially important factor from our results is the size of the calculus deposit. We found a strong positive correlation between biofilm size and retained starch granules (Figure 4A); a result that contradicts findings from archaeological contexts (Dudgeon & Tromp, 2014; Wesolowski et al., 2010). When the concentration of starch granules per mg calculus is considered, the correlation is weaker, but still present (Figure 4B). While the larger deposits contain a higher absolute count, our findings also suggest that they contain a slightly higher concentration of starches. This may also explain the lower mean retention of starch granules in mixed treatments compared to wheat treatments. Wheat treatment samples (mean = 5.53 mg) were on average larger than mixed treatment samples (mean = 4.28 mg) (Table 1); and while mixed treatment solutions contained the highest mean overall granule counts, wheat treatment samples had the highest mean starch retention. Further research is needed to determine why this differs from previous archaeological findings.

Previous research conducted on dental calculus from contemporary humans and non-human primates suggest a high level of stochasticity involved in the retention of starch granules in dental calculus, and that starch granules extracted from dental calculus are underrepresented with regard to actual starch intake, which is consistent with our findings (illustrated by high standard deviations and low proportional incorporation). Leonard and colleagues (2015) found individual calculus samples to be a poor predictor of diet in a population, as many of the consumed plants were missing from some individual samples, but were present in others.

Power and colleagues (2015) presented similar findings in non-human primates, where phytoliths were more representative of individual diets than starch granules. The size bias is also consistent with the findings by Power and colleagues (2015), who found that plants producing starches 10–20 *μ*m in size were over-represented; however, the representation of granules larger than 20 *μ*m in their study is unclear.

The mechanism by which starch granules are incorporated into plaque and calculus remains largely unknown, and few studies have directly investigated potential mechanisms. We know that a proportion of the starch granules entering the mouth can become trapped in the plaque/calculus, and can be recovered from archaeological samples of considerable age (Buckley et al., 2014; Henry et al., 2014; Wu et al., 2021). Studies have also shown that not all starch granules come from a dietary source. Other pathways include cross-contamination from plant interactions in soil, such as palm phytoliths adhering to the skin of sweet potatoes (Tromp & Dudgeon, 2015), or accidental ingestion not related to food consumption (Radini et al., 2017; Radini et al., 2019).

When starch granules enter the mouth, whether through ingestion of food or accidental intake, they immediately encounter multiple obstacles. It is likely that the bulk of starch granules are swallowed along with the food, and are only briefly present in the oral cavity. Other granules that are broken off during mastication may be retained in the dentition through attachment to tooth surfaces (including plaque and dental calculus) and mucous membranes (Dodds & Edgar, 1988; S. Kashket et al., 1991). Bacteria also have the ability to adhere to starch granules (Topping et al., 2003), which would allow starches to attach to bacterial communities within the biofilm. These granules are then susceptible to mechanical removal by the tongue, salivary clearance, and hydrolysis by *α*-amylase (S. Kashket et al., 1996). The susceptibility of granules to hydrolysis depends on the crystallinity and size of the starch granule, as well as the mode of processing. Smaller and pre-processed (e.g., cooked) starch granules are more susceptible to enzymatic degradation, while dehydrated starches will have a reduced susceptibility (Björck et al., 1984; Franco et al., 1992; Haslam, 2004; Henry et al., 2009; Lingstrom et al., 1994). Cracks on the surface of the dental calculus, as well as unmineralised islands and channels may also be able to contain starch granules (Charlier et al., 2010; Power et al., 2014; Tan, Gillam, et al., 2004). Starch granules that are trapped in these pockets are (at least to some extent) protected from aforementioned clearance mechanisms, especially once a new layer of plaque has covered the surface of the plaque/calculus. The size bias against large granules (>20 *μ*m) from both wheat and potato (Table 4) may give further credence to this incorporation pathway, as the smaller starch granules have an advantage over larger granules, and can be stored in larger quantities. This was also suggested by Power and colleagues (2014), who observed clusters of starches within dental calculus, rather than an even distribution across the surface of the dental calculus. Granules trapped in plaque/calculus may still be susceptible to hydrolysis, as *α*-amylase has the ability to bind to both tooth enamel and bacteria within a biofilm and retain a portion of its hydrolytic activity (Nikitkova et al., 2013; Scannapieco et al., 1993; Tan, Mordan, et al., 2004; Tan, Gillam, et al., 2004). After the death of an individual, starches within dental calculus are susceptible to further degradation by post-depositional processes, depending on burial environment (pH, temperature, moisture content, microorganisms) (Franco et al., 1992; García-Granero, 2020; Haslam, 2004; Henry et al., 2009). Future study should explore how burial affects the recovery of starch from the biofilm model.

The absence of *α*-amylase in the model is a limitation of this study, as the total granule counts were not subject to hydrolysis. This would likely have reduced and affected the size ratios, as smaller starches may be more susceptible to hydrolysis (Franco et al., 1992; Haslam, 2004); however, the absence can also allow us to directly explore the effect of *α*-amylase on starch counts in future experiments, where *α*-amylase can be added to the model in concentrations similar to those found in the oral cavity (Scannapieco et al., 1993). We are able to show how absolute counts in the treatments cause a difference in incorporation. However, this was merely a side-effect of the difference in the number of granules in potato and wheat solutions of the same concentration (w/v). Further research should test multiple differing concentrations of the same starch type. The use of EDTA may also have affected counts. While previous studies have shown negligible morphological changes caused by exposure to EDTA (Le Moyne & Crowther, 2021; Modi et al., 2020; Tromp et al., 2017), these studies have not considered changes to separate size categories within starch types, and whether shifts in size ratios occur due to exposure to the pre-treatment chemicals. The total number of granules on a slide often exceeded a number that was feasible to count in a reasonable time period, so we calculated the total counts by extrapolating from three slide transects. Thus, we reasonably assume that the three transects are a good representation of the entire slide, and that the distribution of all granules on the slide is relatively homogeneous.

Finally, we only used native starches in the experimental procedure and the results will likely differ for processed starches (García-Granero, 2020). Based on the comparatively low counts obtained by Leonard and colleagues (2015, Supplement 2), processing and amylase may have a substantial effect on starch granule retention in the oral cavity.

While we are unable to su?iciently address the mechanism(s) of starch incorporation with the data obtained in this study, the dental calculus model presented here is uniquely suited to explore these questions and may improve interpretations of dietary practices in past populations. Further analyses using this model can address the call for more baseline testing of biases associated with dietary research conducted on dental calculus (Le Moyne & Crowther, 2021). Our high-throughput experimental setup allows us a higher degree of control over the factors that influence starch incorporation and retention, such as dietary intake, differential survivability of starches, and inter- and intra-individual variation in plaque accumulation and mineralisation. The latter is especially di?icult to control *in vivo* as it is influenced by numerous factors including genetics, diet, salivary flow, and tooth position and morphology (Fagernäs et al., 2021; Haffajee et al., 2009; Jepsen et al., 2011; Proctor et al., 2018; Simón-Soro et al., 2013), as well as evolutionary differences (Yates et al., 2021). It can also facilitate training of students and researchers on methods of dental calculus analysis, such as starch and phytolith extraction and identification, where it can replace the use of finite archaeological resources.

## 5 Conclusions

This preliminary study shows that a very small proportion of the input starch granules are retained in a dental calculus model. This and previous studies have shown that calculus has a low capacity for retention of starch granules, an effect that is compounded by diagenetic effects in archaeological remains, resulting in low overall counts of extracted granules. The proportion of starches consumed will in many cases be reflected in the quantity of starches extracted from the dental calculus—i.e., the more starch granules entering the oral cavity, the more will be recovered from extraction—at least in modern calculus samples unaffected by diagenesis and hydrolysis. Whether or not this also applies to archaeological samples remains to be tested. Additionally, we have shown that the size of granules will influence the likelihood of incorporation, as large (>20 *μ*m) starches have a decreased incorporation rate, medium (10–20 *μ*m) starches an increased rate, and small (<10 *μ*m) granules remained somewhat constant. The size of calculus deposit also seems to influence the capacity of granule incorporation; as the size of the deposit increases, so does the absolute count of incorporated granules.

While we have shown multiple factors that influence the likelihood of incorporation, the process still appears to be somewhat stochastic. Further research is needed to make sense of the contributing factors, and to explore the mechanisms of intra-oral starch incorporation and retention in dental calculus. The oral biofilm model described in this study provides a method to explore the incorporation and extraction of dietary compounds from dental calculus in a controlled laboratory setting, and unearth the potential biases associated with dietary research conducted on archaeological dental calculus.

## Acknowledgements

We would like to thank Dr. Stephanie Schnorr for help with the amylase activity protocol. We also thank everyone in the general vicinity of the lab for enduring the smell of bacterial accumulation. Since we did NOT make use of Sci-Hub to access articles stuck behind a paywall, we will NOT acknowledge the use of Sci-Hub in this study.

This research has received funding from the European Research Council under the European Union’s Horizon 2020 research and innovation program, grant agreement number STG–677576 (“HARVEST”).

## Data Availability Statement

All scripts and data used in the analysis are available on OSF (https://osf.io/uc5qy/) and Github (https://github.com/bbartholdy/byoc-starch), following the format provided by the rrtools package (Marwick, 2019). More detailed protocols are available on OSF (https://osf.io/akevs/) and protocols.io (https://www.protocols.io/view/biofilm-growth-with-starch-treatment-bu7jnzkn). Additional tables and figures are available in the Supplementary Material.

