## Supplementary Material for "Investigating Biases Associated with Dietary Starch Incorporation and Retention with an Oral Biofilm Model"

### R Session info

```
print(sessionInfo(), locale = FALSE)
```

```
## R version 4.1.1 (2021-08-10)
## Platform: x86_64-pc-linux-gnu (64-bit)
## Running under: Pop!_OS 21.04
##
## Matrix products: default
## BLAS:   /usr/lib/x86_64-linux-gnu/blas/libblas.so.3.9.0
## LAPACK: /usr/lib/x86_64-linux-gnu/lapack/liblapack.so.3.9.0
##
## attached base packages:
## [1] stats     graphics  grDevices utils     datasets  methods   base
##
## other attached packages:
##  [1] byocstarch_0.0.0.9000 broom_0.7.7           patchwork_1.1.1
##  [4] forcats_0.5.1         stringr_1.4.0         dplyr_1.0.6
##  [7] purrr_0.3.4           readr_1.4.0           tidyr_1.1.3
## [10] tibble_3.1.4          ggplot2_3.3.4         tidyverse_1.3.1
## [13] here_1.0.1
##
## loaded via a namespace (and not attached):
##  [1] Rcpp_1.0.7        lubridate_1.7.10  prettyunits_1.1.1 ps_1.6.0
##  [5] assertthat_0.2.1  rprojroot_2.0.2   digest_0.6.28     utf8_1.2.2
```

```
##  [9] R6_2.5.1          cellranger_1.1.0  backports_1.2.1   reprex_2.0.0
## [13] evaluate_0.14     httr_1.4.2        pillar_1.6.3      rlang_0.4.11
## [17] readxl_1.3.1      rstudioapi_0.13   callr_3.7.0       rmarkdown_2.9
## [21] desc_1.4.0        devtools_2.4.2    munsell_0.5.0     compiler_4.1.1
## [25] modelr_0.1.8      xfun_0.26         pkgconfig_2.0.3   pkgbuild_1.2.0
## [29] htmltools_0.5.2   tidyselect_1.1.1  bookdown_0.23     fansi_0.5.0
## [33] crayon_1.4.1      dbplyr_2.1.1      withr_2.4.2       grid_4.1.1
## [37] jsonlite_1.7.2    gtable_0.3.0      lifecycle_1.0.1   DBI_1.1.1
## [41] magrittr_2.0.1    scales_1.1.1      cli_3.0.1         stringi_1.7.4
## [45] cachem_1.0.6      remotes_2.4.0     fs_1.5.0          testthat_3.0.4
## [49] xml2_1.3.2        ellipsis_0.3.2    generics_0.1.0    vctrs_0.3.8
## [53] tools_4.1.1       glue_1.4.2        hms_1.1.0         pkgload_1.2.2
## [57] processx_3.5.2    fastmap_1.1.0     yaml_2.2.1        colorspace_2.0-1
## [61] sessioninfo_1.1.1 rvest_1.0.0       memoise_2.0.0     knitr_1.36
## [65] haven_2.4.1       usethis_2.0.1
```

### Metadata for raw data files

Counts represent the absolute number of starches counted on a slide

**starch_counts.csv**

| variable | description |
|---|---|
| sample | Sample number. |
| plate | Plate number that the sample came from. |
| row | Which row on the plate the sample came from. |
| s | Small starch count. |
| m | Medium starch count. |
| l | Large starch count. |
| total | Sum of s, m, and l. |
| treatment | Treatment solution to which the samples were exposed. |
| starch | Type of starch that was counted. |
| weight | Weight of the biofilm sample. |
| vol | Total volume of EDTA in which the sample was dissolved. |
| portion_slide | Proportion of the microscope slide that was counted. Total transects on slide divided by counted transects. |

**solution_counts.csv**

| variable | description |
|---|---|
| solution | Type of starch in solution. |
| concentration | Concentration (%w/v) of starch in solution. |
| vol_slide | Volume of solution added to slide. |
| vol_total | Total volume of solution in aliquot. |
| portion_slide | Proportion of slide that was counted. Total transects on slide divided by counted transects. |
| slide | Slide number. |
| starch | Starch type counted. |
| s | Small starch count. |

| variable | description |
|---|---|
| m | Medium starch count. |
| l | Large starch count. |
| total | Sum of s, m, and l. |

### Raw data

The raw data can be downloaded from GitHub:

```
# solution counts
wget https://github.com/bbartholdy/byoc-starch/blob/main/analysis/data/raw_data/solution_counts.csv

# sample counts
wget https://github.com/bbartholdy/byoc-starch/blob/main/analysis/data/raw_data/starch_counts.csv
```

Raw counts from the treatment solutions before extrapolation.

| solution | concentration | vol_slide | vol_total | portion_slide | slide | starch | s | m | l | total |
|---|---|---|---|---|---|---|---|---|---|---|
| wheat | 0.25 | 10 | 1000 | 0.1034483 | 1 | wheat | 969 | 387 | 167 | 1523 |
| wheat | 0.25 | 10 | 1000 | 0.1034483 | 2 | wheat | 1118 | 445 | 199 | 1762 |
| potato | 0.25 | 10 | 1000 | 0.1034483 | 1 | potato | 9 | 95 | 86 | 190 |
| potato | 0.25 | 10 | 1000 | 0.1034483 | 2 | potato | 7 | 78 | 115 | 200 |
| mix | 0.25 | 10 | 1000 | 0.1034483 | 1 | wheat | 1218 | 414 | 116 | 1748 |
| mix | 0.25 | 10 | 1000 | 0.1034483 | 1 | potato | NA | 68 | 60 | 128 |

Raw counts from the calculus samples before extrapolation:

| sample | plate | row | s | m | l | total | treatment | starch | weight | vol | portion_slide |
|---|---|---|---|---|---|---|---|---|---|---|---|
| st1A1 | 1 | A | 39 | 535 | 119 | 693 | potato | potato | 5.80 | 100 | 1.0000000 |
| st1A2 | 1 | A | 6 | 28 | 5 | 39 | potato | potato | 5.81 | 100 | 1.0000000 |
| st1A3 | 1 | A | 26 | 1392 | 389 | 1807 | potato | potato | 8.22 | 100 | 1.0000000 |
| st1A4 | 1 | A | 14 | 184 | 40 | 238 | potato | potato | 4.65 | 100 | 1.0000000 |
| st1A5 | 1 | A | 20 | 341 | 98 | 459 | potato | potato | 7.68 | 200 | 1.0000000 |
| st1A6 | 1 | A | 32 | 466 | 159 | 657 | potato | potato | 7.79 | 200 | 1.0000000 |
| st1B1 | 1 | B | 62 | 36 | 4 | 102 | wheat | wheat | 5.15 | 100 | 0.1034483 |
| st1B2 | 1 | B | 508 | 321 | 18 | 847 | wheat | wheat | 4.56 | 100 | 0.2500000 |
| st1B3 | 1 | B | 606 | 664 | 73 | 1343 | wheat | wheat | 9.28 | 100 | 0.1034483 |
| st1B4 | 1 | B | 61 | 51 | 14 | 126 | wheat | wheat | 1.59 | 100 | 0.1034483 |
| st1B5 | 1 | B | 276 | 227 | 64 | 567 | wheat | wheat | 8.62 | 200 | 0.1034483 |
| st1B6 | 1 | B | 175 | 96 | 19 | 290 | wheat | wheat | 9.80 | 200 | 0.1034483 |
| st1C1 | 1 | C | NA | 57 | 19 | 76 | mix | potato | 4.09 | 100 | 0.1034483 |
| st1C1 | 1 | C | 97 | 94 | 50 | 241 | mix | wheat | 4.09 | 100 | 0.1034483 |
| st1C2 | 1 | C | NA | 12 | 13 | 25 | mix | potato | 1.50 | 100 | 0.1034483 |
| st1C2 | 1 | C | 31 | 30 | 9 | 70 | mix | wheat | 1.50 | 100 | 0.1034483 |
| st1C3 | 1 | C | NA | 113 | 20 | 133 | mix | potato | 8.44 | 100 | 0.1034483 |
| st1C3 | 1 | C | 351 | 256 | 39 | 646 | mix | wheat | 8.44 | 100 | 0.1034483 |
| st1C4 | 1 | C | NA | 78 | 25 | 103 | mix | potato | 5.42 | 100 | 0.1034483 |
| st1C4 | 1 | C | 392 | 302 | 68 | 762 | mix | wheat | 5.42 | 100 | 0.1034483 |
| st1C5 | 1 | C | NA | 22 | 10 | 32 | mix | potato | 6.12 | 200 | 1.0000000 |

| sample | plate | row | s | m | l | total | treatment | starch | weight | vol | portion_slide |
|---|---|---|---|---|---|---|---|---|---|---|---|
| st1C5 | 1 | C | 5 | 0 | 0 | 5 | mix | wheat | 6.12 | 200 | 1.0000000 |
| st1C6 | 1 | C | NA | 17 | 0 | 17 | mix | potato | 1.91 | 100 | 1.0000000 |
| st1C6 | 1 | C | 97 | 52 | 12 | 161 | mix | wheat | 1.91 | 100 | 1.0000000 |
| st1D1 | 1 | D | NA | NA | NA | 1 | control | none | 6.51 | 100 | 1.0000000 |
| st1D2 | 1 | D | NA | NA | NA | 0 | control | none | 4.42 | 100 | 1.0000000 |
| st1D3 | 1 | D | NA | NA | NA | 0 | control | none | 5.01 | 200 | 1.0000000 |
| st1D4 | 1 | D | NA | NA | NA | 0 | control | none | 5.14 | 100 | 1.0000000 |
| st1D5 | 1 | D | NA | NA | NA | 0 | control | none | 4.51 | 100 | 1.0000000 |
| st1D6 | 1 | D | NA | NA | NA | 0 | control | none | 1.67 | NA | NA |
| st2A1 | 2 | A | 20 | 150 | 24 | 194 | potato | potato | 6.11 | 200 | 1.0000000 |
| st2A2 | 2 | A | 89 | 479 | 34 | 602 | potato | potato | 2.54 | 100 | 1.0000000 |
| st2A3 | 2 | A | 71 | 370 | 22 | 463 | potato | potato | 8.48 | 200 | 1.0000000 |
| st2A4 | 2 | A | 59 | 773 | 135 | 967 | potato | potato | 5.91 | 200 | 1.0000000 |
| st2A5 | 2 | A | 97 | 512 | 292 | 901 | potato | potato | 8.92 | 200 | 1.0000000 |
| st2A6 | 2 | A | NA | NA | NA | NA | potato | potato | 3.14 | NA | NA |
| st2B1 | 2 | B | 183 | 130 | 20 | 333 | wheat | wheat | 8.08 | 200 | 0.1034483 |
| st2B2 | 2 | B | 27 | 19 | 3 | 49 | wheat | wheat | 2.30 | 100 | 0.1034483 |
| st2B3 | 2 | B | 585 | 409 | 43 | 660 | wheat | wheat | 6.84 | 100 | 0.1034483 |
| st2B4 | 2 | B | 32 | 21 | 2 | 55 | wheat | wheat | 0.56 | 100 | 0.1034483 |
| st2B5 | 2 | B | 308 | 263 | 46 | 617 | wheat | wheat | 8.51 | 200 | 0.1034483 |
| st2B6 | 2 | B | NA | NA | NA | NA | wheat | wheat | 1.06 | NA | NA |
| st2C1 | 2 | C | NA | 79 | 17 | 96 | mix | potato | 5.04 | 100 | 0.1034483 |
| st2C1 | 2 | C | 521 | 331 | 58 | 910 | mix | wheat | 5.04 | 100 | 0.1034483 |
| st2C2 | 2 | C | NA | 25 | 1 | 26 | mix | potato | 3.64 | 100 | 0.1034483 |
| st2C2 | 2 | C | 182 | 101 | 25 | 308 | mix | wheat | 3.64 | 100 | 0.1034483 |
| st2C3 | 2 | C | NA | 31 | 4 | 35 | mix | potato | 4.11 | 100 | 0.1034483 |
| st2C3 | 2 | C | 252 | 142 | 19 | 413 | mix | wheat | 4.11 | 100 | 0.1034483 |
| st2C4 | 2 | C | NA | 43 | 13 | 56 | mix | potato | 3.61 | 100 | 0.1034480 |
| st2C4 | 2 | C | 327 | 222 | 45 | 594 | mix | wheat | 3.61 | 100 | 0.1034480 |
| st2C5 | 2 | C | NA | 14 | 0 | 14 | mix | potato | 3.17 | 100 | 1.0000000 |
| st2C5 | 2 | C | 14 | 8 | 0 | 22 | mix | wheat | 3.17 | 100 | 1.0000000 |
| st2C6 | 2 | C | NA | NA | NA | NA | mix | potato | 1.75 | NA | NA |
| st2D1 | 2 | D | 0 | 0 | 0 | 0 | control | none | 8.32 | 100 | 1.0000000 |
| st2D2 | 2 | D | 0 | 0 | 0 | 0 | control | none | 11.18 | 200 | 1.0000000 |
| st2D3 | 2 | D | NA | NA | NA | NA | control | none | 3.43 | NA | NA |
| st2D4 | 2 | D | NA | NA | NA | NA | control | none | 5.76 | NA | NA |
| st2D5 | 2 | D | NA | NA | NA | NA | control | none | 3.66 | NA | NA |
| st2D6 | 2 | D | NA | NA | NA | NA | control | none | 5.67 | NA | NA |

### Experimental setup

```
knitr::include_graphics(here("analysis/figures/plate_lid_side.jpg"))
```

```
knitr::include_graphics(here("analysis/figures/plate_lid_on.jpg"))
```

### Protocols

All protocols are available on protocols.io.

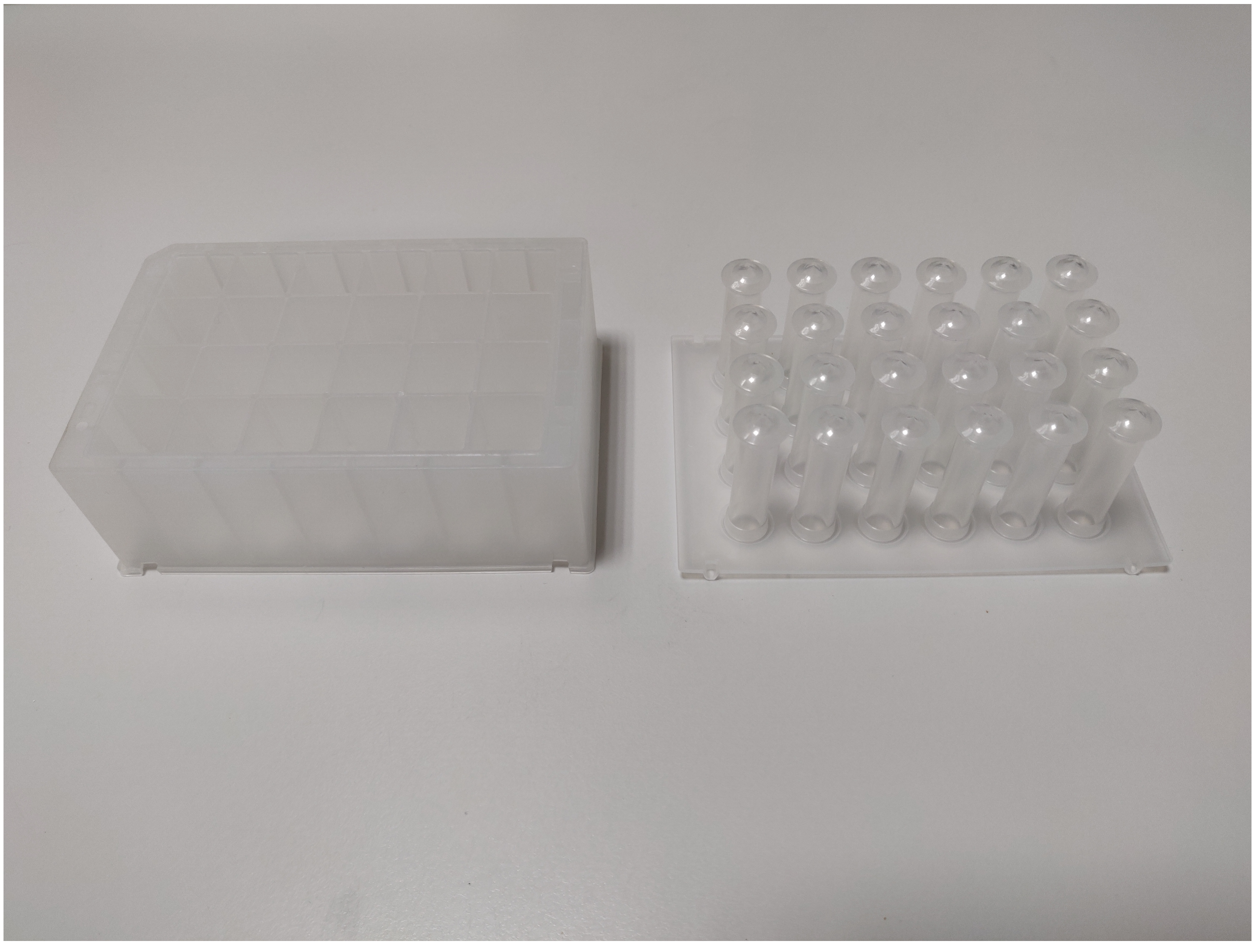

Figure 1: The 24 deepwell plate and the lid with pegs (substrata)

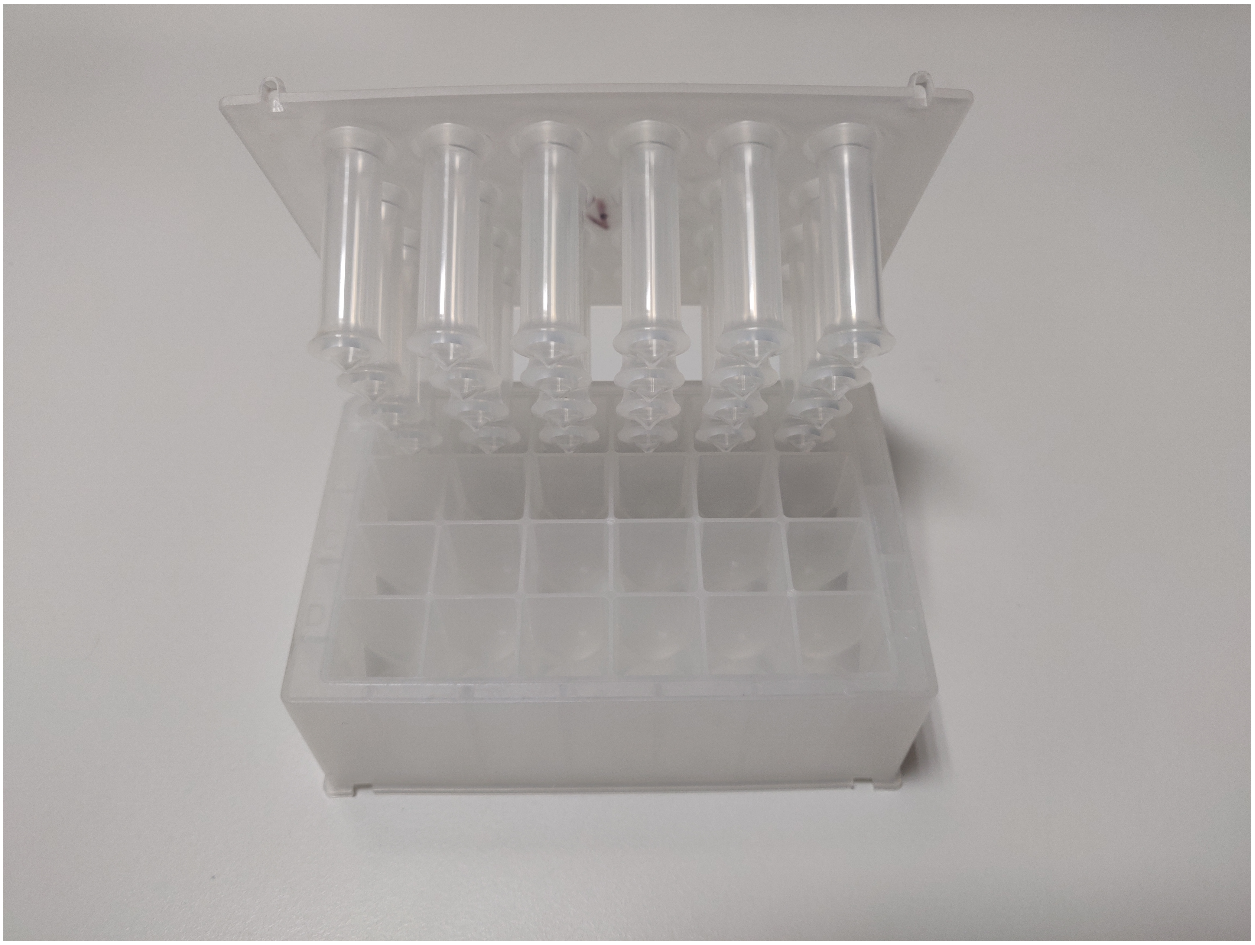

Figure 2: The 24 deepwell plate with the lid (almost) on.

Creating the artificial saliva: https://www.protocols.io/view/artificial-saliva-bva9n2h6

Creating the CPMU solution: https://www.protocols.io/view/cpmu-bv8pn9vn

Biofilm growth protocol: https://www.protocols.io/view/biofilm-growth-with-starch-treatment-bu7jnzkn

Amylase activity assay: https://www.protocols.io/view/amylase-activity-bw8jphun

### Microscope images

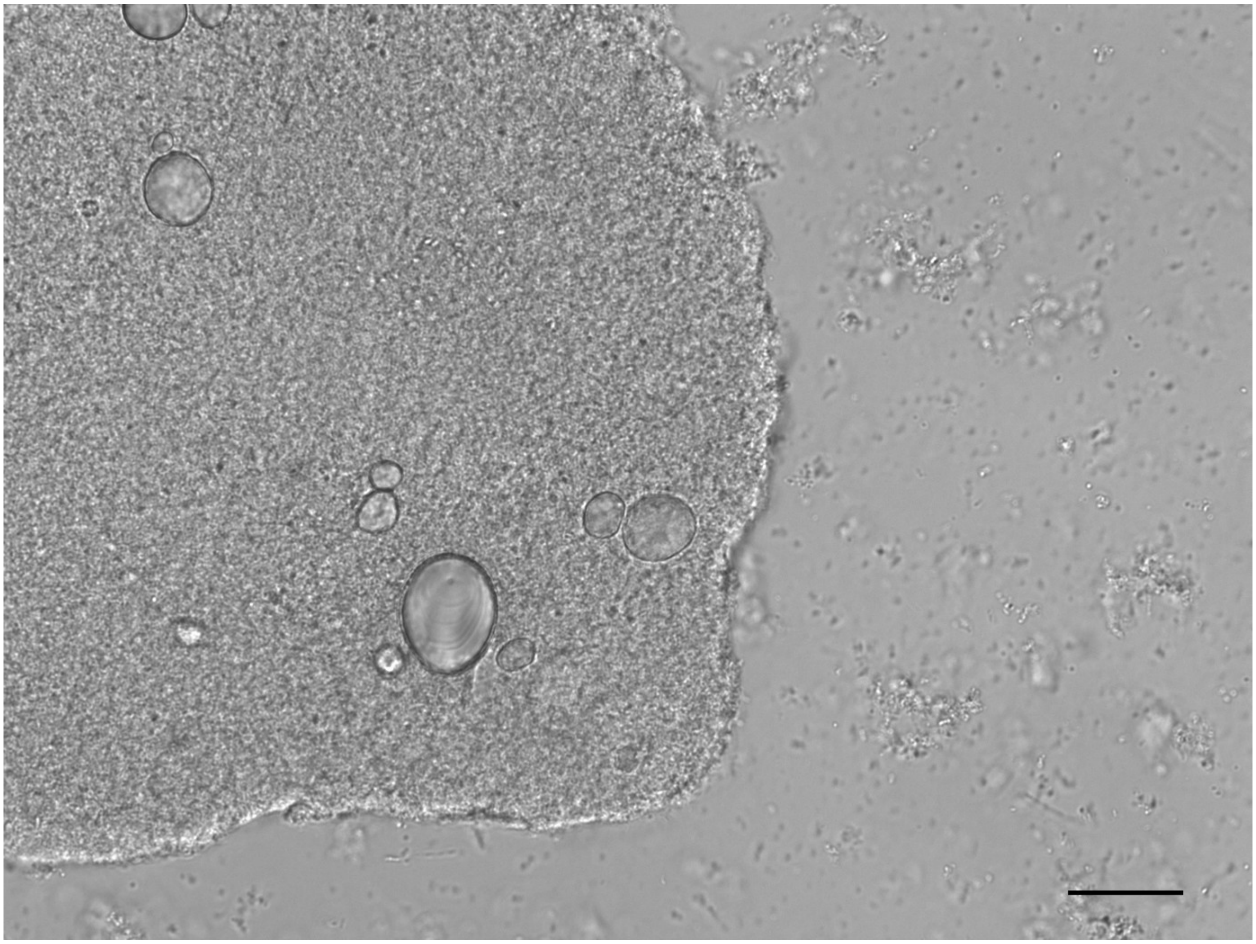

Figure 3: Image of starch granules extracted from a potato treatment sample

```
knitr::include_graphics(here("analysis/figures/SNAP-103412-0006.jpg"))
```

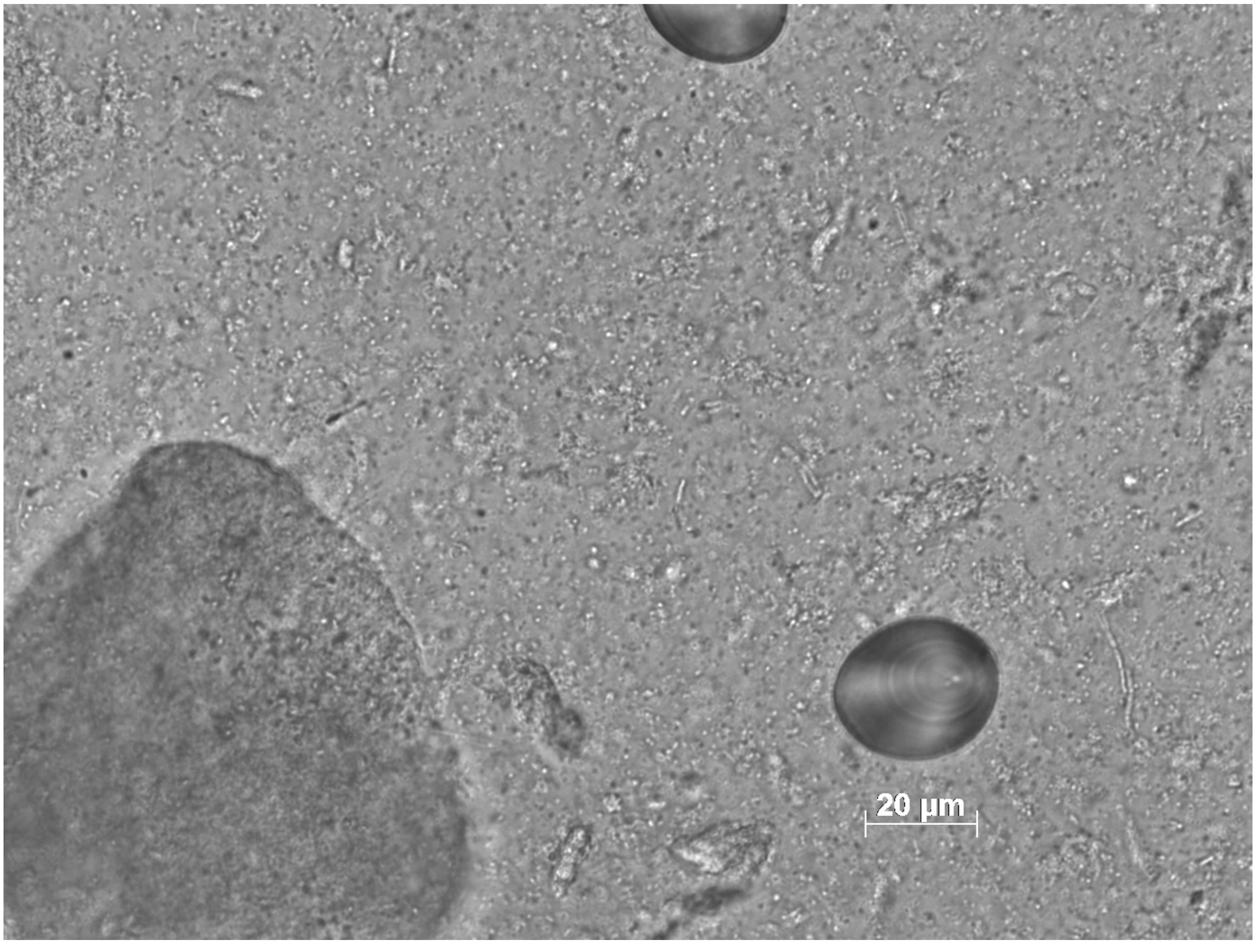

```
knitr::include_graphics(here("analysis/figures/SNAP-164650-0012.jpg"))
```

### Amylase activity

Amylase activity in U/mL enzyme, where U is mg maltose released from starch in six minutes at 36 °C.

Tables containing the amylase activity results for both plates and both photometric readings conducted on each plate. Samples (rows) were analysed in triplicates (columns).

```
# table of results reported in units amylase per mL enzyme (but let's be honest,
  # ...it doesn't really matter what the unit is. No activity is no activity)
cols <- c("1", "2", "3")  # sample triplicates
rows <- c("S1", "S2", "S3", "B1", "B2", "B3", "B4", "B5", "BT1", "BT2", "BT3")
plt1_ph1_result <- rbind(sal1_ph1, bmm1_ph1)
rownames(plt1_ph1_result) <- rows
plt1_ph2_result <- rbind(sal1_ph2, bmm1_ph2)
rownames(plt1_ph2_result) <- rows
plt2_ph1_result <- rbind(sal2_ph1, bmm2_ph1)
rownames(plt2_ph1_result) <- rows
plt2_ph2_result <- rbind(sal2_ph2, bmm2_ph2)
rownames(plt2_ph2_result) <- rows
```

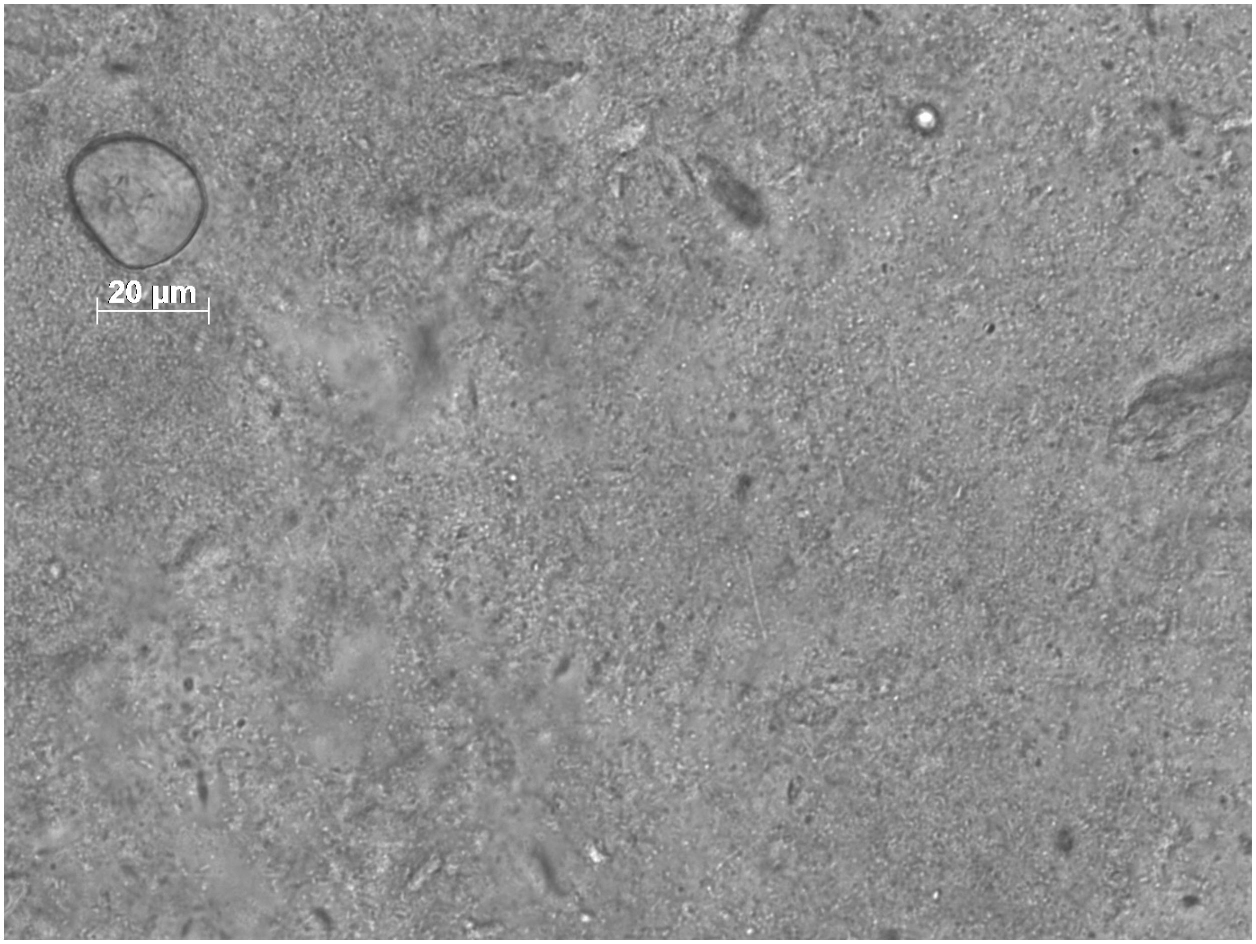

Figure 4: Microscope image of wheat starch from a wheat treatment sample.

```
plt1_ph1_result # plate 1, photometric reading 1
```

| | V1 | V2 | V3 |
|---|---|---|---|
| S1 | 9.6633544 | 3.4437165 | 9.7409060 |
| S2 | 10.2992774 | 4.7465833 | 9.6090682 |
| S3 | 9.1902896 | 5.1498516 | 9.6711095 |
| B1 | -0.2944638 | -0.2420948 | -0.2682793 |
| B2 | -0.1940899 | -0.3642891 | -0.2464589 |
| B3 | -0.2115462 | -0.4210222 | -0.1504490 |
| B4 | -0.2726434 | -0.4384786 | -0.3119201 |
| B5 | -0.3381046 | -0.3599251 | -0.2202744 |
| BT1 | -0.4952116 | -0.4384786 | -0.4864835 |
| BT2 | -0.4864835 | -0.3031920 | -0.4952116 |
| BT3 | -0.5083039 | -0.4341145 | -0.4690271 |

```
plt1_ph2_result # plate 1, photometric reading 2
```

| | V1 | V2 | V3 |
|---|---|---|---|
| S1 | 9.5791455 | 3.3993902 | 9.5869680 |
| S2 | 10.2049435 | 4.7292109 | 9.6260804 |
| S3 | 9.1567319 | 5.1516246 | 9.7199501 |
| B1 | -0.2739597 | -0.2345669 | -0.2520748 |
| B2 | -0.1689122 | -0.3527452 | -0.2345669 |
| B3 | -0.1864201 | -0.4096459 | -0.1426504 |
| B4 | -0.2476978 | -0.4271538 | -0.2958445 |
| B5 | -0.3221064 | -0.3527452 | -0.2170590 |
| BT1 | -0.4796775 | -0.4271538 | -0.4753006 |
| BT2 | -0.4709236 | -0.2914676 | -0.4796775 |
| BT3 | -0.4928085 | -0.4227768 | -0.4534157 |

```
plt2_ph1_result # plate 2, photometric reading 1
```

| | V1 | V2 | V3 |
|---|---|---|---|
| S1 | 9.6074482 | 3.5463151 | 9.2241068 |
| S2 | 10.3307337 | 4.6674077 | 9.4989553 |
| S3 | 8.9854226 | 5.2677348 | 9.5351196 |
| B1 | -0.2451714 | -0.2745476 | -0.2745476 |
| B2 | -0.1990089 | -0.3920521 | -0.2619578 |
| B3 | -0.1780260 | -0.4675907 | -0.1864192 |
| B4 | -0.2997271 | -0.4717873 | -0.3374964 |
| B5 | -0.2577612 | -0.4004453 | -0.2325817 |
| BT1 | -0.5011634 | -0.4088385 | -0.4759839 |
| BT2 | -0.5011634 | -0.2913339 | -0.4885737 |
| BT3 | -0.5305396 | -0.3291032 | -0.5179498 |

```
plt2_ph2_result # plate 2, photometric reading 2
```

| | V1 | V2 | V3 |
|---|---|---|---|
| S1 | 9.6115786 | 3.5595447 | 9.1398973 |
| S2 | 10.2719324 | 4.6335267 | 9.3721096 |
| S3 | 8.9149416 | 5.2430841 | 9.4301627 |
| B1 | -0.2266208 | -0.2644543 | -0.2602506 |
| B2 | -0.1887873 | -0.3779548 | -0.2434357 |
| B3 | -0.1635650 | -0.4578255 | -0.1677687 |
| B4 | -0.2896767 | -0.4578255 | -0.3191027 |
| B5 | -0.2476394 | -0.3905660 | -0.2182134 |
| BT1 | -0.4956590 | -0.3947697 | -0.4662330 |
| BT2 | -0.4914553 | -0.2728618 | -0.4746404 |
| BT3 | -0.5166776 | -0.3106953 | -0.5040665 |

### Control samples

```
raw_counts %>%
  filter(treatment == "control") %>%
  select(!c(vol, portion_slide, s, m, l))
```

| sample | plate | row | total | treatment | starch | weight |
|---|---|---|---|---|---|---|
| st1D1 | 1 | D | 1 | control | none | 6.51 |
| st1D2 | 1 | D | 0 | control | none | 4.42 |
| st1D3 | 1 | D | 0 | control | none | 5.01 |
| st1D4 | 1 | D | 0 | control | none | 5.14 |
| st1D5 | 1 | D | 0 | control | none | 4.51 |
| st1D6 | 1 | D | 0 | control | none | 1.67 |
| st2D1 | 2 | D | 0 | control | none | 8.32 |
| st2D2 | 2 | D | 0 | control | none | 11.18 |
| st2D3 | 2 | D | NA | control | none | 3.43 |
| st2D4 | 2 | D | NA | control | none | 5.76 |
| st2D5 | 2 | D | NA | control | none | 3.66 |
| st2D6 | 2 | D | NA | control | none | 5.67 |

Only the total starch count was considered for control samples, as size was deemed irrelevant.

### Count corrections

Slide transects were calculated by counting the number of transects on the cover slip under the microscope. This was done by starting in the bottom-left corner, and counting the total number of full fields-of-view across the cover slip to the bottom-right corner. The total number of transects was 29 (verified multiple times).

A 1 mL aliquot of each of the original treatment solutions was taken, from which 10 $\mu$L was taken and mounted on a microscope slide and mixed with 10 $\mu$L 20% (v/v) glycerol. Solution counts were extrapolated from a slide (10 $\mu$L) to the quantity in a 1 ml solution, and then multipled by 16 days to achieve the total number of granules that were exposed to the samples:

$$\text{corrected count} = \text{raw count} \times \frac{\text{total slides}}{\text{counted slides}} \times 100\mu L \times 16 \text{ days}$$

Samples were submerged in 50–100 $\mu$L EDTA, from which 20 $\mu$L was mounted on a microscope slide ($V_{slide}$) and counted. Sample counts were extrapolated to the full volume of EDTA ($V_{sample}$) in which the sample was submerged (i.e. 50–100 $\mu$L).

$$\text{Corrected count} = \text{raw count} \times (\text{portion of slide})^{-1} \times \frac{V_{sample}}{V_{slide}}$$

#### Some additional plots

Bar plot for the total count of granules exposed to the samples over the duration of the experiment,

```
sol_long %>%
  filter(size == "total") %>%
  group_by(treatment, starch) %>%
  ggplot(aes(x = treatment, y = count, fill = treatment, col = starch)) +
  geom_col(size = 1.5) +
  theme(panel.background = element_rect(fill = "white"),
        panel.grid = element_line(colour = "grey"),
        panel.grid.major.x = element_blank(),
        axis.title.x = element_blank()) +
  scale_fill_viridis_d() +
  scale_color_viridis_d(begin = 0.5)
```

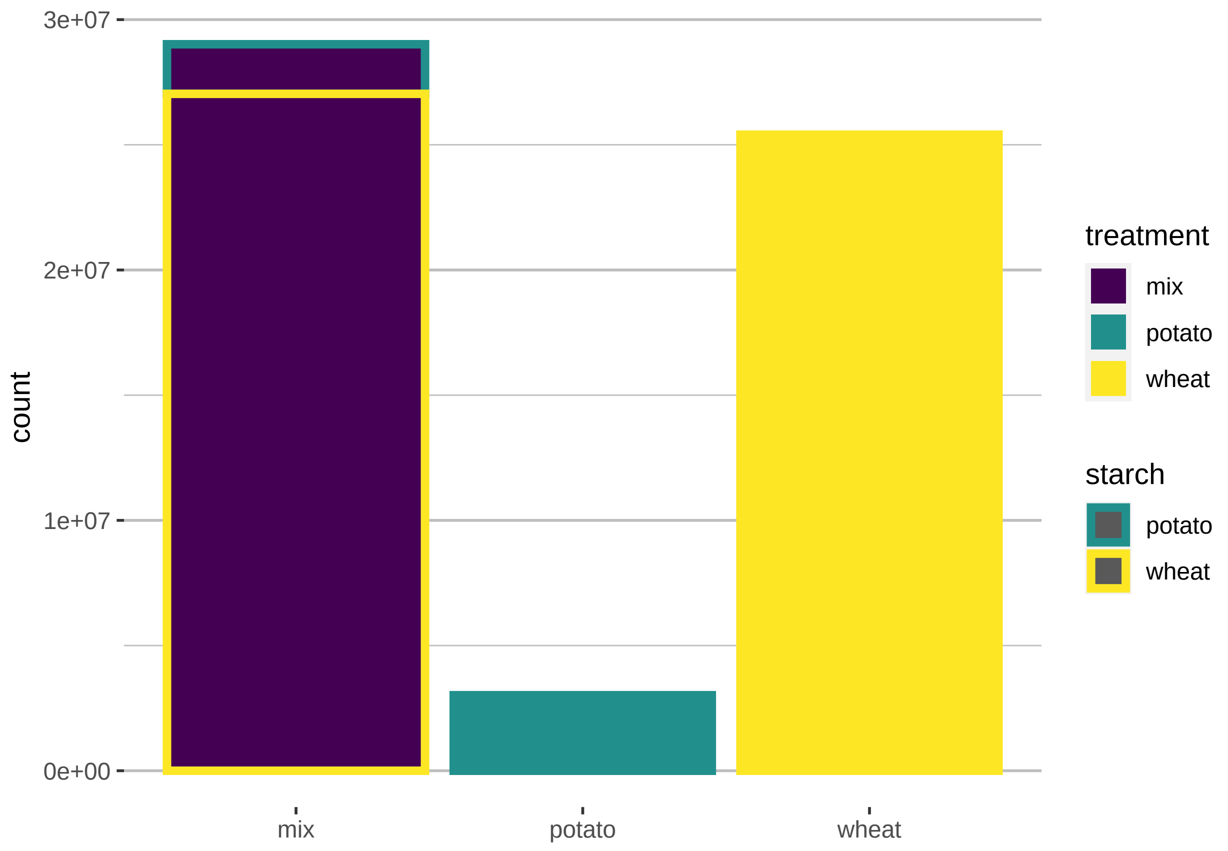

and box plot with superimposed points (with added jitter) for the extrapolated mean counts of granules extracted from the samples.

```r
corr_comb %>%
  filter(treatment != "control") %>%
  ggplot(aes(x = treatment, y = total,
             shape = treatment)) +
    geom_boxplot(aes(fill = treatment), alpha = 0.5) +
    geom_jitter(aes(col = treatment), width = 0.3, size = 2) +
    scale_color_viridis_d() +
    theme(panel.background = element_rect(fill = "white"),
        panel.grid = element_line(colour = "grey"),
        panel.grid.major.x = element_blank(),
        axis.title.x = element_blank()) + # remove y-axis title
    scale_fill_viridis_d()
```

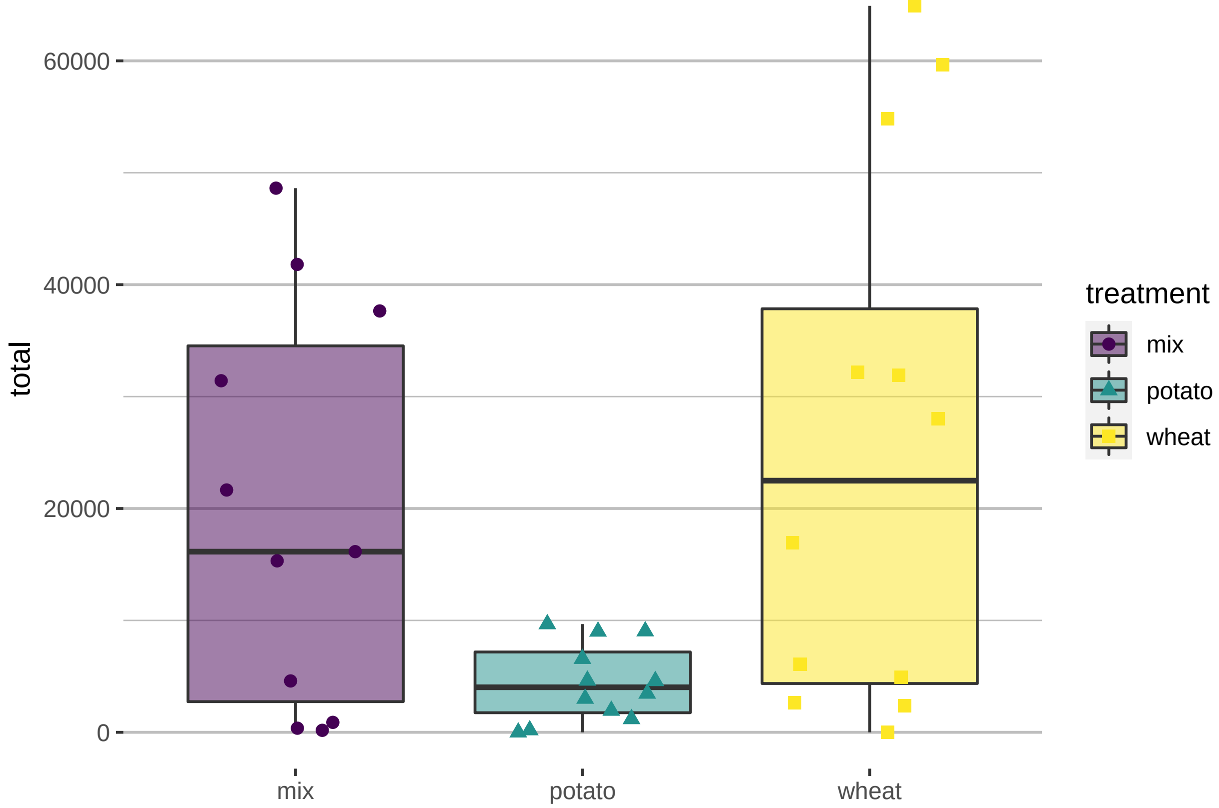

Extracted-granule counts separated by treatment and size, including error bars:

```r
corr_counts_long %>%
  filter(size != "total",
         treatment != "control") %>%
  group_by(treatment, starch, size) %>%
  summarise(sd = sd(count, na.rm = T),
            count = mean(count, na.rm = T)) %>%
  #mutate(percent = count / sum(count, na.rm = T) * 100) %>%
  ggplot(aes(x = starch, y = count, fill = size)) +
    geom_col(position = "dodge") +
    geom_errorbar(aes(ymin = count, ymax = count + sd), width = 0.2, position = position_dodge(0.9)) +
    facet_wrap(~ treatment, scales = "free") +
```

```
    scale_fill_viridis_d() +
    theme_bw()
```

```
## 'summarise()' has grouped output by 'treatment', 'starch'. You can override using the '.groups' argu

## Warning: Removed 1 rows containing missing values (geom_col).
```

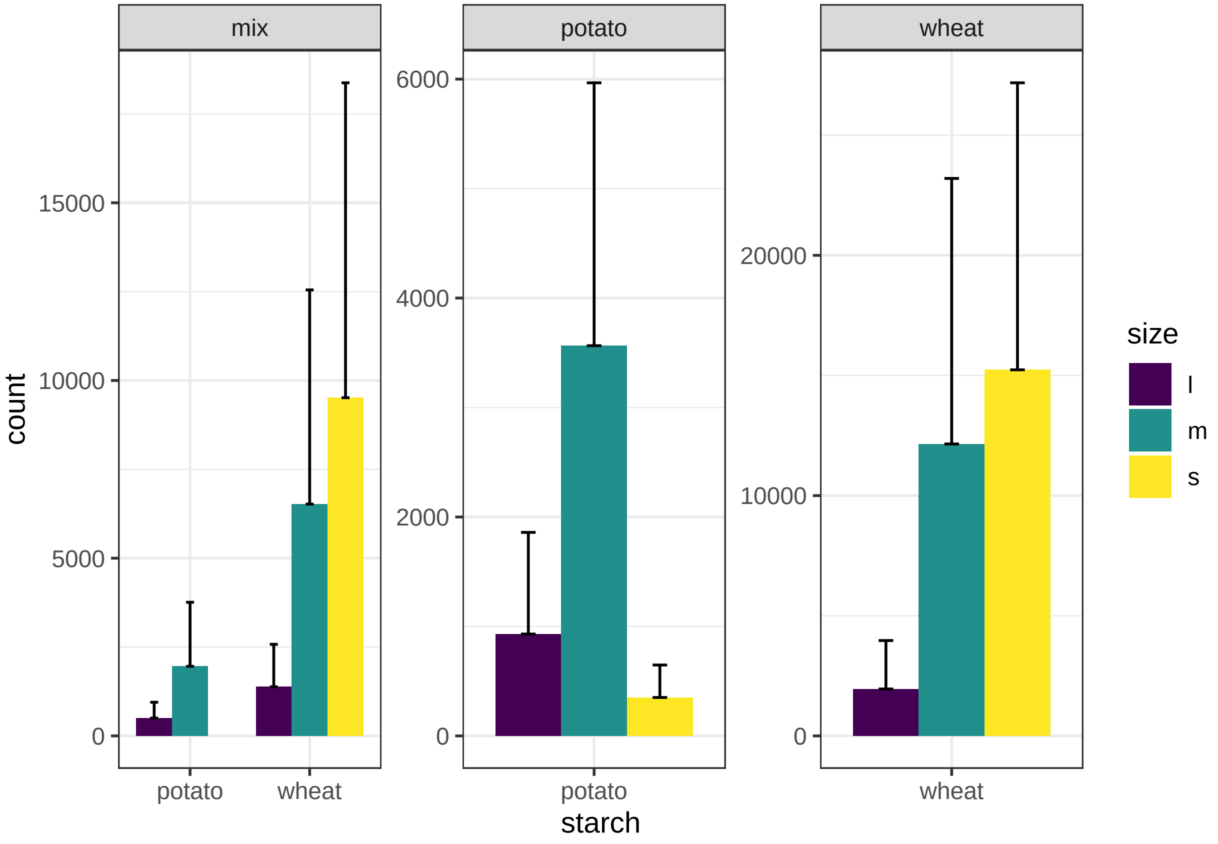

Figure 5: l = large, m = medium, s = small.

Size distribution (in %) within the solutions (top) and samples (bottom):

```
sol_size_pl <- sol_corr %>%
  group_by(solution, starch) %>%
  summarise(across(c(s, m, l, total), mean, na.rm = T)) %>%
  pivot_longer(cols = c(s,m,l, total), values_to = "count", names_to = "size") %>%
  filter(size != "total") %>%
  group_by(solution, starch) %>%
  mutate(percent = count / sum(count, na.rm = T) * 100) %>%
  ggplot(aes(x = starch, y = percent, fill = size)) +
    geom_col(position = "dodge") +
    facet_wrap(~ solution, scales = "free_x") +
    scale_fill_viridis_d() +
    theme_bw() +
    labs(x = "")
```

```r
samp_size_pl <- corr_counts_long %>%
  filter(size != "total",
         treatment != "control") %>%
  group_by(treatment, starch, size) %>%
  summarise(count = mean(count, na.rm = T)) %>%
  mutate(percent = count / sum(count, na.rm = T) * 100) %>%
  ggplot(aes(x = starch, y = percent, fill = size)) +
    geom_col(position = "dodge") +
    facet_wrap(~ treatment, scales = "free_x") +
    scale_fill_viridis_d() +
    theme_bw()

sol_size_pl / samp_size_pl + plot_layout(guides = "collect")
```

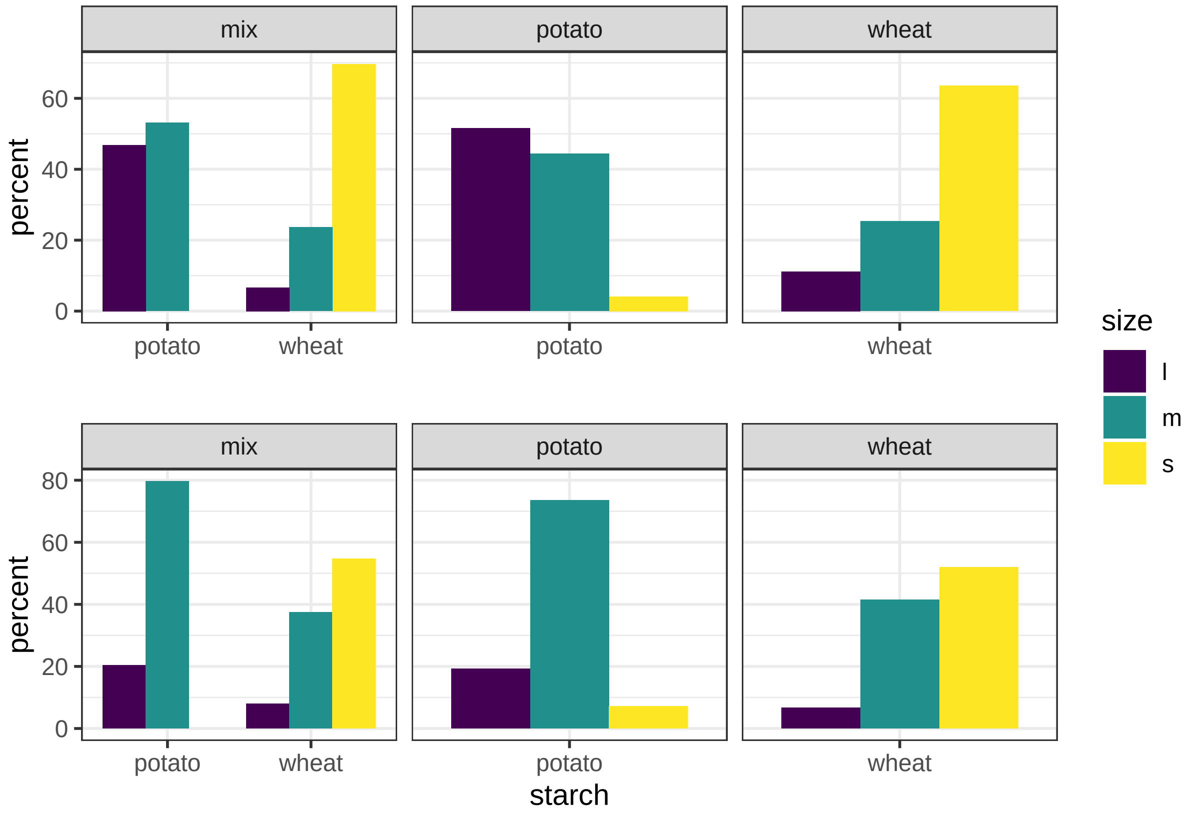

Figure 6: l = large, m = medium, s = small.

Separated correlation plots. These are the same plots as in the main paper, just larger.

```r
pl_cor
```

```r
pl_cor2
```

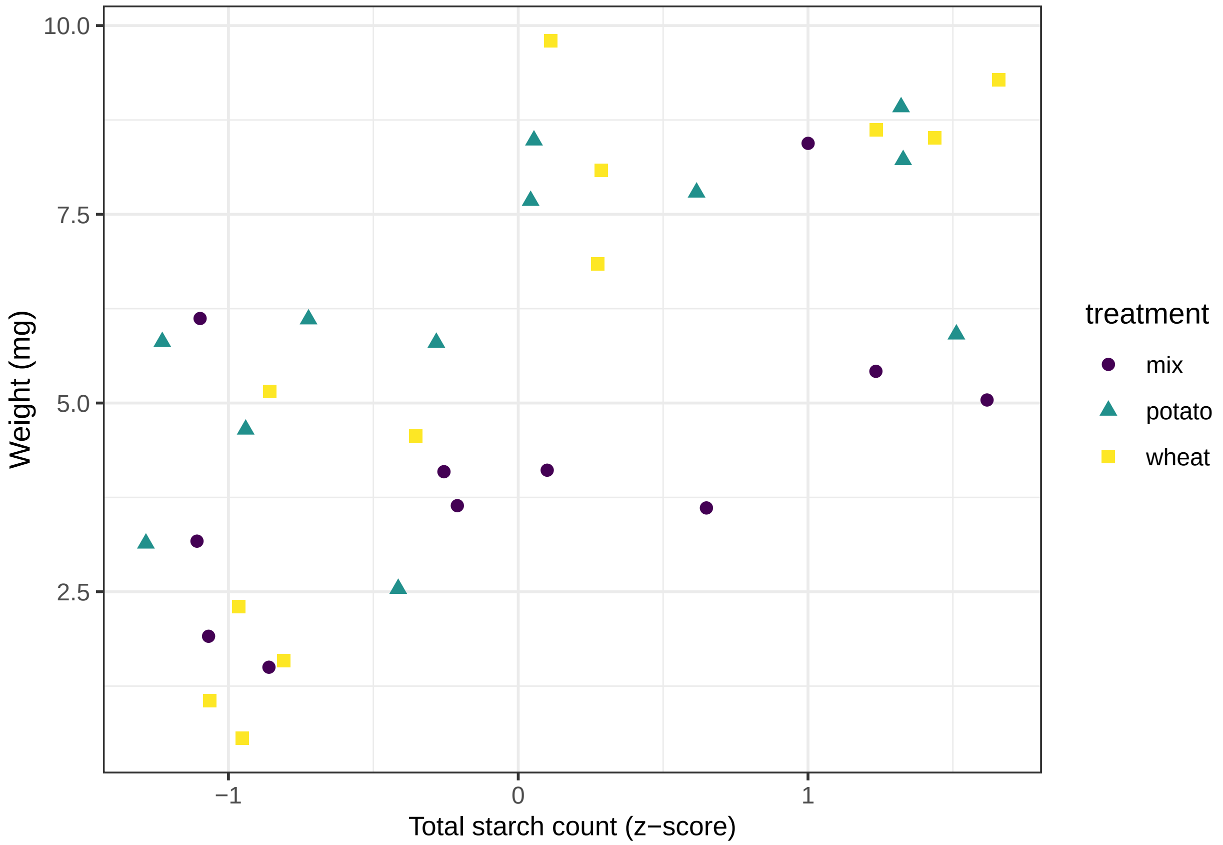

Figure 7: Scatter plot of sample weight and standardised starch count by z-score for separated treatments.

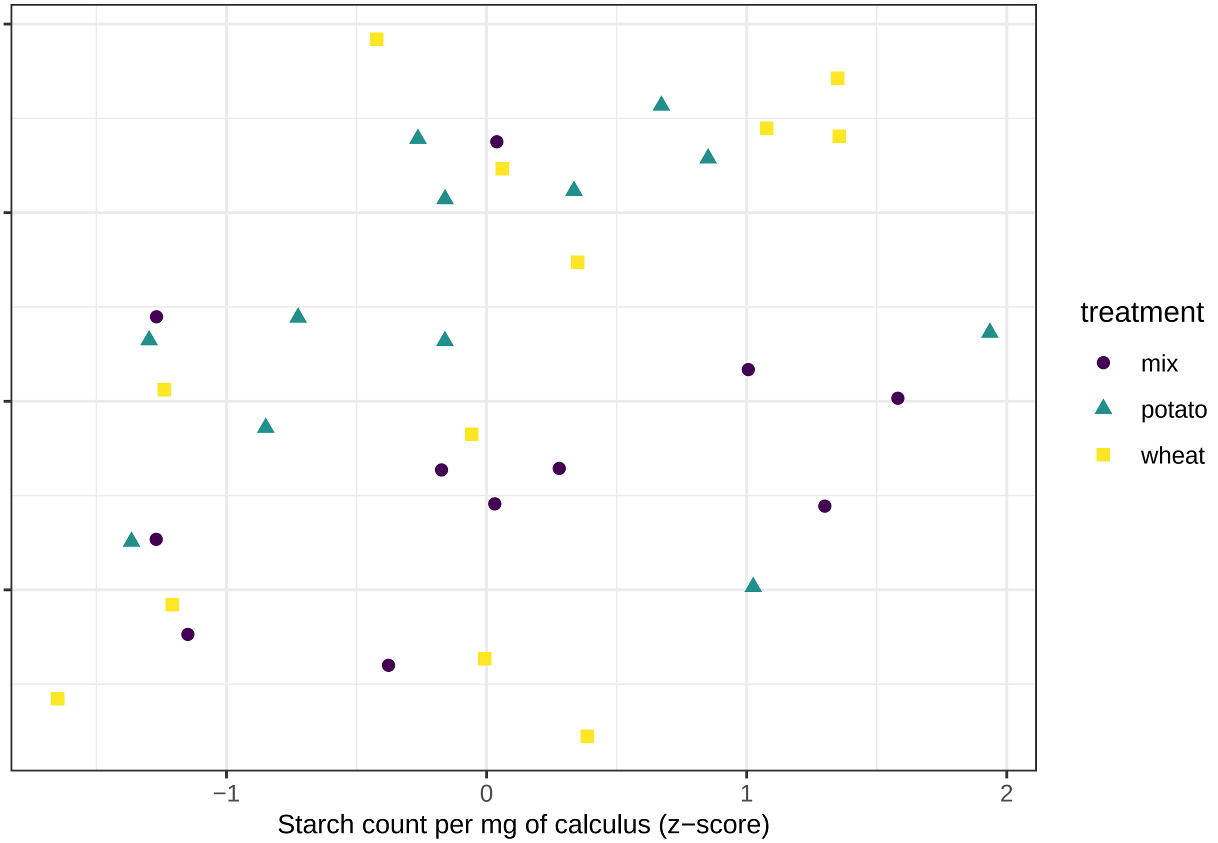

Figure 8: Scatter plot of sample weight in mg and standardised count of starch grains per mg calculus.

#### ...and a table

Differences in size ratios (%) of granules between the solutions and the samples. Negative values indicate a loss of granules from solution to sample.

```
size_diff %>%
  mutate(across(where(is.numeric), signif, 3))
```

| treatment | starch | s | m | l |
|---|---|---|---|---|
| mix | potato | NaN | 26.5 | -26.5000 |
| mix | wheat | -15.00 | 13.8 | 1.2900 |
| mix | both | -17.10 | 17.0 | 0.0863 |
| potato | potato | 3.14 | 29.2 | -32.3000 |
| wheat | wheat | -8.49 | 18.6 | -4.0900 |
